# Crystal structure and catalytic mechanism of PL35 family glycosaminoglycan lyases with an ultrabroad substrate spectrum

**DOI:** 10.1101/2024.09.04.611182

**Authors:** Lin Wei, Hai-Yan Cao, Ruyi Zou, Min Du, Qingdong Zhang, Danrong Lu, Xiangyu Xu, Yingying Xu, Wenshuang Wang, Xiu-Lan Chen, Yu-Zhong Zhang, Fuchuan Li

## Abstract

Recently, a new class of glycosaminoglycan (GAG) lyases (GAGases) belonging to PL35 family has been discovered with an ultrabroad substrate spectrum that can degrade three types of uronic acid-containing GAGs (hyaluronic acid, chondroitin sulfate and heparan sulfate) or even alginate. In this study, the structures of GAGase II from *Spirosoma fluviale* and GAGase VII from *Bacteroides intestinalis* DSM 17393 were determined at 1.9 and 2.4 Å resolution, respectively, and their catalytic mechanism was investigated by the site-directed mutant of their crucial residues and molecular docking assay. Structural analysis showed that GAGase II and GAGase VII consist of an N-terminal (α/α)_6_ toroid multidomain and a C-terminal two-layered β-sheet domain with Mn^2+^. Notably, although GAGases share similar folds and catalytic mechanisms with some GAG lyases and alginate lyases, they exhibit higher structural similarity with alginate lyases than GAG lyases, which may present a crucial structural evidence for the speculation that GAG lyases with (α/α)_n_ toroid and antiparallel β-sheet structures arrived by a divergent evolution from alginate lyases with the same folds. Overall, this study not only solved the structure of PL35 GAG lyases for the first time and investigated their catalytic mechanism, especially the reason why GAGase III can additionally degrade alginate, but also provided a key clue in the divergent evolution of GAG lyases that originated from alginate lyases.

## Introduction

Glycosaminoglycans (GAGs) are a class of linear polyanionic polysaccharides ubiquitously distributed on the cell surface and in the extracellular matrix (ECM) of animal tissues (***Cohen and Merzendorfer, 2019***), and they participate in various physiological and pathological processes through interacting with a series of chemokines (***Dyer, et al., 2015***; ***Irie, et al., 2008***), growth factors (***Sirko, et al., 2010***; ***Zhang, et al., 2019***) or other ECM components. Except for keratan sulfate (KS), which does not contain hexuronic acid (HexUA) residues, other GAGs are composed of repeating disaccharides consisted of HexUA (D-glucuronic acid (GlcUA) or L-iduronic acid (IdoUA)) and hexosamine (HexNAc) (D-*N*-acetylgalactosamine (GalNAc) or D-*N*- acetylglucosamine (GlcNAc)). Based on the disaccharide composition and the type of glycosidic bonds between disaccharides units, HexUA-containing GAGs are classified into three classes: hyaluronan (HA), chondroitin sulfate (CS)/dermatan sulfate (DS) and heparan sulfate (HS)/heparin (Hep) (***Cohen and Merzendorfer, 2019***). HA is the only non-sulfated GAG composed of repeating -GlcUAβ1-3GlcNAc- disaccharides linked by β1-4 glycosidic bonds. In contrast, the structures of CS/DS and Hep/HS are highly complex due to sulfation and epimerization modifications. The backbone of CS is composed of repeating -GlcUAβ1- 3GalNAc- disaccharides linked by β1-4 glycosidic bonds, and is further sulfated at C-2 of GlcUA and C-4/C-6 of GalNAc by sulfotransferases; meanwhile, the GlcUA residues in CS can be epimerized into IdoUA residues by glucuronyl C-5 epimerase to form DS (***Kusche-Gullberg and Kjellen, 2003***). Similarly, HS/Hep composed of repeating -4GlcUAβ1-4GlcNAcα1- units can be sulfated at C-2 of GlcUA and N/C-2/C-3/C-6 of GlcNAc/deacetylated GlcN, and GlcUA residues are often epimerized to IdoUA residues, especially in Hep (***Cohen and Merzendorfer, 2019***). Such complex modification through sulfation and epimerization endows sulfated GAGs with diverse biological functions.

GAG lyases, belonging to the superfamily of polysaccharide lyases (PLs), specifically catalyze the degradation of HexUA-containing GAGs (***Charnock, et al., 2002***; ***Garron and Cygler, 2010***). These reactions generate a plethora of oligosaccharides that contain an unsaturated double bond between C-4 and C-5 on uronic acids at their nonreducing ends, which is generated via a β-elimination mechanism (***Garron and Cygler, 2010***). As a complimentary mechanistic strategy to glycoside hydrolases (GHs), GAG lyases degrade HexUA-containing GAGs without the participation of a water molecule and involved in the polysaccharide metabolism of microorganisms (***Garron and Cygler, 2010***; ***Ndeh, et al., 2020***). For the past many years, an increasing number of gene sequences containing PL catalytic modules and some ancillary modules have been annotated as PLs (***Lombard, et al., 2010***). Based on their sequence similarity, 44 families (and an unclassified PL0 family) have been hierarchically classified in the CAZy database (http://www.cazy.org/), and at least one member of each family has been subjected to activity identification and biochemical property analysis (***Drula, et al., 2022***). The identified GAG lyases are widely distributed in the PL6, PL8, PL12, PL13, PL15, PL16, PL21, PL23, PL29, PL30, PL33, PL35 and PL37 families (***Drula, et al., 2022***). Based on substrate specificity, GAG lyases are classified as HA-specific lyases, which can degrade HA only, CS/DS lyases (chondroitinase, CSases), which degrade CS/DS as well as HA in most cases, and Hep/HS lyases (heparinases, Hepases), which specifically cleave Hep/HS. All identified GAG lyases have substrate specificity to strictly distinguish one or two kinds of GAGs based on saccharide composition and glycosidic bonds between each HexUA-HexNAc disaccharide units, while other uronic acid-containing polysaccharides are usually unable to serve as substrates for these enzymes.

With the evolution of host polysaccharides (***Csoka and Stern, 2013***; ***Popper, et al., 2011***), the bacterial enzymes that degrade these polysaccharides have also evolved to varying degrees in response to complex environments. As a kind of unbranched anionic extracellular polysaccharide produced by lower organisms algae and bacteria (such as brown alga (***Popper, et al., 2011***) and bacteria belonging to *Pseudomonas* (***Evans and Linker, 1973***) and *Azotobacter* (***Clementi, 1997***), alginate containing no hexosamine and sulfation possesses a simpler structure than GAGs in animals and likely emerge earlier than GAGs during the course of evolution. Like GAGs, alginate is a linear polyanionic polysaccharide that contains HexUA residues; however, alginate possesses a completely different chemical structure, which is composed of D-mannuronate (M) and its C5 epimer L-guluronate (G), and M and G residues often alternate randomly to form heteropolyuronic blocks, including polyM composed of M residues linked by β1-4 bonds, polyG composed of G residues linked by α1-4 bonds, and polyMG/GM composed of alternating M and G linked by β/α1-4 bonds (***Pawar and Edgar, 2012***). Likewise, a series of alginate lyases essential for the metabolism of alginate were identified from microorganisms and algae as well as lower marine animals, which belong to the PL5, PL6, PL7, PL8, PL14, PL15, PL17, PL18, PL31, PL32, PL34, PL36, PL38, PL39, PL41 and PL44 families (***Drula, et al., 2022***). Based on their specificity, these lyases are also divided into the following categories: M block-specific lyases (***Itoh, et al., 2019***; ***Zhu, et al., 2015***), G block-specific lyases (***Matsubara, et al., 1998***; ***Yamasaki, et al., 2005***) and MG-specific alginate lyases (***Jagtap, et al., 2014***; ***Yamasaki, et al., 2004***). From an evolutionary perspective in previous reports, GAG lyases are thought to have originated from alginate lyases by a divergent evolution through adaptation to the evolution of substrate polysaccharides. Some similarities in the sequences and folds of these two types of enzymes support this idea. For example, the (α/α)_n_ toroid and antiparallel β-sheet folds of GAG lyases are found in families such as PL8 (hyaluronate lyase and chondroitin lyase), PL12 (heparinase III), PL21 (heparinase II), and PL23 (chondroitin lyase). Alginate lyases in PL15, PL17, and PL39 families also exhibit these folds. Additionally, the β-jelly roll fold of heparinase I in the PL13 family and alginate lyases in PL7 and PL18 families show similar structures. Lastly, the β-helix fold of chondroitinase B and alginate lyase in the PL6 family is another example (***Garron and Cygler, 2014***). Considering that the alginate with simpler structure from brown alga or bacteria might appear earlier, it is possible that the ancestral enzymes originally used alginate as a substrate. However, enzymes with transitional characteristics in activity and structure have not been discovered to support this speculation.

Recently, a new class of GAG lyases belonging to PL35 family was discovered with an ultrabroad substrate spectrum (***Wei, et al., 2024***). Unlike the identified GAG lyases (HA-specific lyases, CSases and Hepases) with quite strict substrate specificities, these novel GAG lyases can degrade three types of uronic acid-containing GAGs (HA, CS and HS), and thus were named GAGases. More interestingly, one of the eight identified GAGases (GAGase III) can even degrade alginate. Analysis of substrate structure preference represented by GAGase I showed that GAGases selectively act on GAG domains composed of non/6-*O*-/*N*-sulfated hexosamines and D-glucuronic acid, among which GAGase III can also selectively degrade polyM blocks composed of D-mannuronic acids but not polyG blocks composed of L-guluronic acids in alginate. Furthermore, the primary structure alignment and phylogenetic analysis showed that GAGases have considerable degree of similarity with not only GAG lyases from PL15, PL21 and PL33 families but also alginate lyases from PL15, PL17 and PL34 families (***Wei, et al., 2024***), indicating that GAGases may exhibit some transitional features from alginate lyases to GAG lyases in structure.

In this study, crystal structures of two GAGases (GAGase II and GAGase VII) were determined for the first time, and their catalytic mechanism was further elucidated through the site-directed mutagenesis of their crucial site residues and molecular docking. Structural alignment showed that GAGases are more structural similarity to some alginate lyases rather than GAG lyases, suggesting that PL35 family GAGases might originate from alginate lyases possessing an N-terminal (α/α)_n_ toroid domain and a C-terminal antiparallel β-sheet domain. Moreover, one of the reasons why GAGase III can additionally degrade M-block alginate are also resolved by structural modeling, structural alignment and site-directed mutagenesis. This study not only helps to resolve the catalytic mechanism of the PL35 family lyases in particular GAGases, but also provides potential structural evidence for the divergent evolution of GAG lyases.

## Results

### Overall crystal structures of GAGase II and GAGase VII

To explore the substrate recognition and catalytic mechanism of the GAGases, the eight enzymes with different sequence identity (*Supplemental Table S1*) were individually used to prepare their crystals for structural analysis. The structures of GAGase II from *Spirosoma fluviale* (GenBank accession number: SOD82962.1, 81.2% sequence identity with GAGase I), GAGase VII from *Bacteroides intestinalis* (GenBank accession number: EDV05210.1, 44.8% sequence identity with GAGase I) and selenomethionine (SeMet)-labelled GAGase II (Se- GAGase II) in their ligand-free states were solved at 1.9 Å, 2.4 Å and 2.0 Å resolution, respectively (*Table 1*). Both GAGase II and GAGase VII are monomers consisting of two domains: a N-terminal (α/α)_6_ toroid domain and a C-terminal two-layered β-sheet domain (*Figure 1A*).

**Figure 1.**
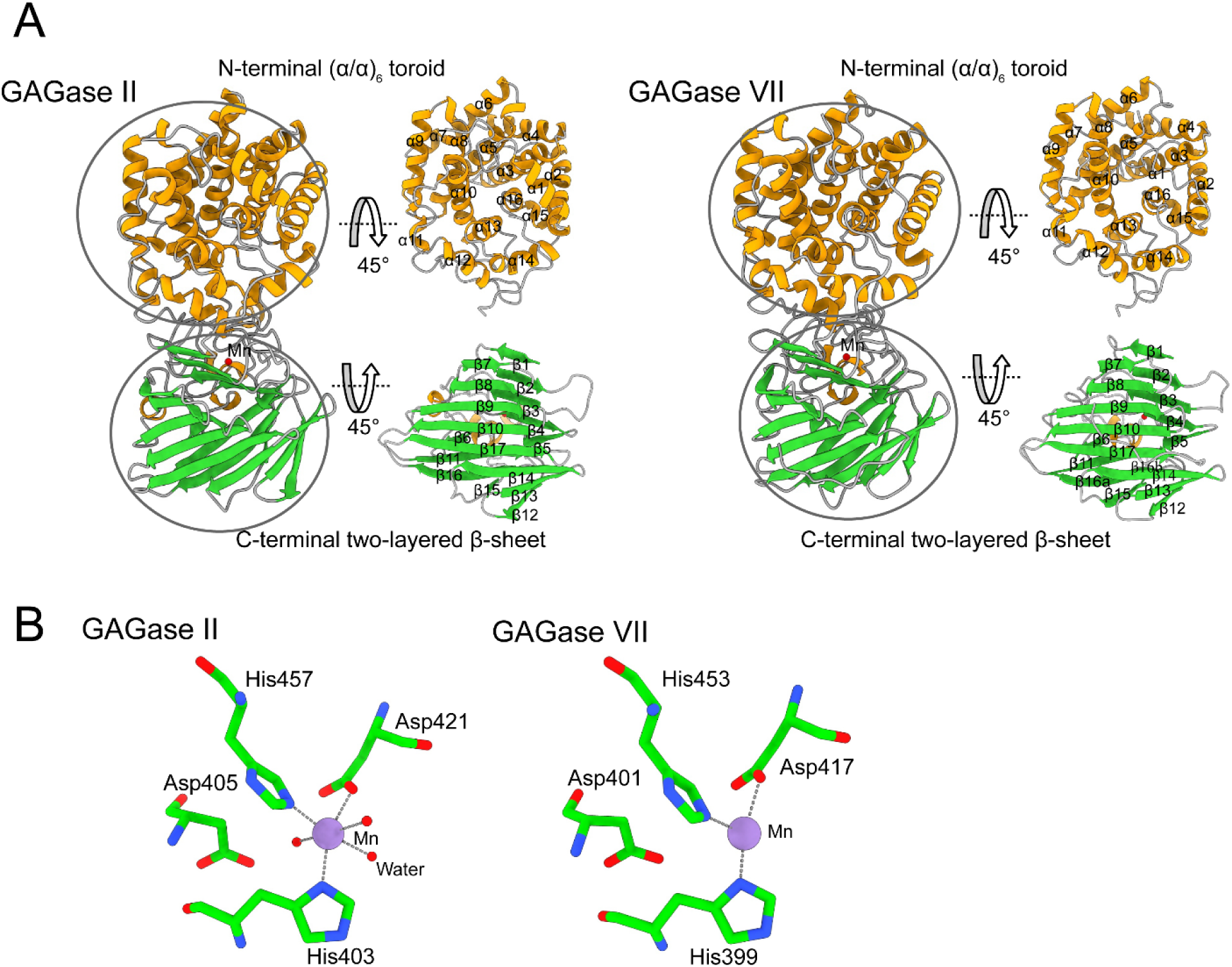
Structural description of GAGase II and GAGase VII. **(A)**, Overall structures of GAGase II (*left*) and GAGase VII (*right*) are shown in carton. The α-helix, β-strand and random coil are colored with yellow, green and gray, respectively. The N-terminal (α/α)_6_ toroid domain and C-terminal two-layered β- sheet domain were circled and the secondary structure elements are marked nearby. **(B)**, Mn binding site of GAGase II and GAGase VII. The *purple* sphere presents Mn^2+^. Mn^2+^ binding site of GAGase II (*left*) and GAGase VII (*right*) is shown in stick. A cut off distance of 3.0 Å was carried out to choose neighboring residues.

**Table 1.**
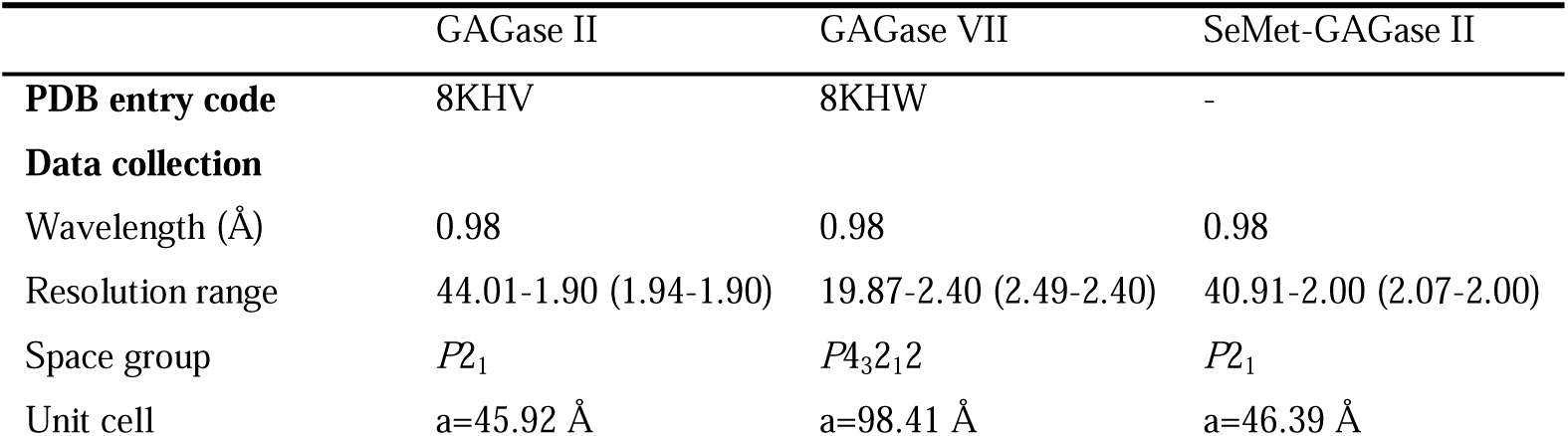
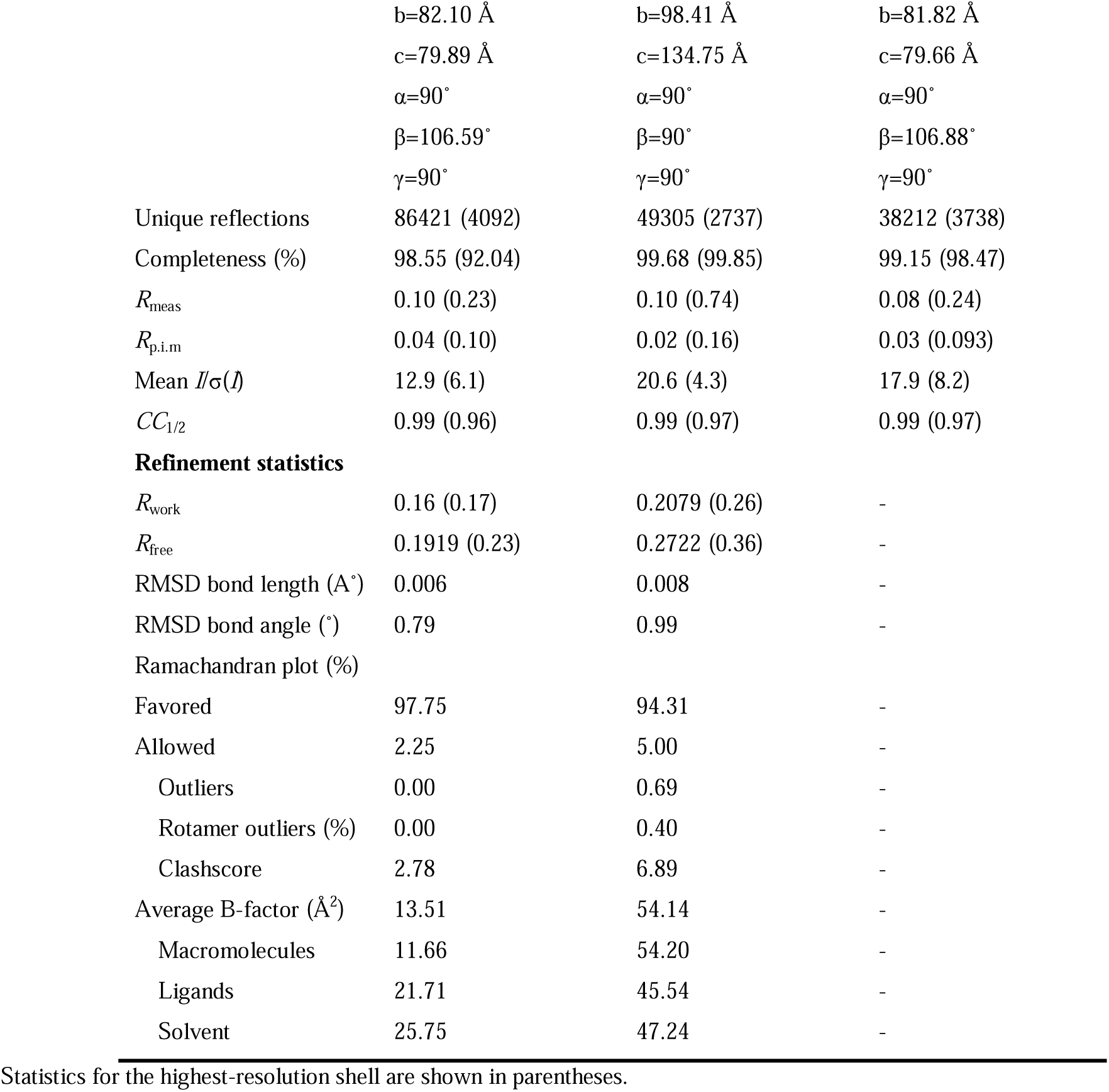
Data collection and refinement statistics.

|  | GAGase II | GAGase VII | SeMet-GAGase II |
| --- | --- | --- | --- |
| **PDB entry code** | 8KHV | 8KHW | - |
| **Data collection** |  |  |  |
| Wavelength (Å) | 0.98 | 0.98 | 0.98 |
| Resolution range | 44.01-1.90 (1.94-1.90) | 19.87-2.40 (2.49-2.40) | 40.91-2.00 (2.07-2.00) |
| Space group | $P2_1$ | $P4_32_12$ | $P2_1$ |
| Unit cell | a=45.92 Å | a=98.41 Å | a=46.39 Å |
|  | b=82.10 Å | b=98.41 Å | b=81.82 Å |
|  | c=79.89 Å | c=134.75 Å | c=79.66 Å |
| | $\alpha=90^\circ$ | $\alpha=90^\circ$ | $\alpha=90^\circ$ |
| | $\beta=106.59^\circ$ | $\beta=90^\circ$ | $\beta=106.88^\circ$ |
| | $\gamma=90^\circ$ | $\gamma=90^\circ$ | $\gamma=90^\circ$ |
| Unique reflections | 86421 (4092) | 49305 (2737) | 38212 (3738) |
| Completeness (%) | 98.55 (92.04) | 99.68 (99.85) | 99.15 (98.47) |
| $R_{\text{meas}}$ | 0.10 (0.23) | 0.10 (0.74) | 0.08 (0.24) |
| $R_{\text{p.i.m}}$ | 0.04 (0.10) | 0.02 (0.16) | 0.03 (0.093) |
| Mean $I/\sigma(I)$ | 12.9 (6.1) | 20.6 (4.3) | 17.9 (8.2) |
| $CC_{1/2}$ | 0.99 (0.96) | 0.99 (0.97) | 0.99 (0.97) |
| **Refinement statistics** |  |  |  |
| $R_{\text{work}}$ | 0.16 (0.17) | 0.2079 (0.26) | - |
| $R_{\text{free}}$ | 0.1919 (0.23) | 0.2722 (0.36) | - |
| RMSD bond length (Å) | 0.006 | 0.008 | - |
| RMSD bond angle (°) | 0.79 | 0.99 | - |
| Ramachandran plot (%) |  |  |  |
| Favored | 97.75 | 94.31 | - |
| Allowed | 2.25 | 5.00 | - |
| Outliers | 0.00 | 0.69 | - |
| Rotamer outliers (%) | 0.00 | 0.40 | - |
| Clashscore | 2.78 | 6.89 | - |
| Average B-factor (Å<sup>2</sup>) | 13.51 | 54.14 | - |
| Macromolecules | 11.66 | 54.20 | - |
| Ligands | 21.71 | 45.54 | - |
| Solvent | 25.75 | 47.24 | - |

Statistics for the highest-resolution shell are shown in parentheses.

The N-terminal domain (Met^34^-Ala^365^) of GAGase II is composed of 16 large or small α- helices (α1-α16) to form an (α/α)_6_ toroid domain. Specifically, the N-terminal 16 α-helices constitute seven complete α-helix pairs (α1-2, α3-4, α5-7, α8-9, α10-12, α13-14 and α15-16), forming seven hairpin structures, which eventually embrace into the (α/α)_6_ toroid structure. The transition between α6 to α8 and between α10 to 12 is discontinuous, and both transitions possess a small α-helix α7 and α11, respectively. In each hairpin structure, two antiparallel α-helices are connected by loops composed of 9-12 residues, and each hairpin contains a turn structure composed of 1-4 residues. The N-terminal of this toroid domain is connected to a signal peptide that has been removed during cloning expression, and its C-terminal is connected to the two-layered β-sheet domain through a 17-residue linkage (Trp^350^-Met^365^). Its inner layer is formed by helices α1, α3, α5, α8, α10, α13 and α16, while its outer layer is formed by α2, α4, α6+α7, α9, α11+α12, α14 and α15. Moreover, the α-helix pairs α13-14 and α15-16 are significantly shorter than the first five sets, and α15 and α16 are connected by a short β-turn consisting of only 3 residues, which makes the toroid structure closed (*Figure 1A*).

The C-terminal domain (Met^366^-Ser^612^) of GAGase II is composed of 17 β-strands (β1-β17) to form a two-layered β-sheet domain with 4 small α-helices located between β5-β6 and β8-β9. Both layers of this C-terminal β-sheet domain are composed of eight groups of β-strands, namely β1-β5, β11 and β15-β16 and β7-β10, β12-β14 and β17. The two layers are roughly parallel to each other and twisted to form an angle of approximately 60°. The first five β-strands (β1-β5) of the C-terminal domain are arranged in anti-parallel configuration. Subsequently, there is an Ω- loop structure wrapping around a metal ion (Mn^2+^), which consists of three small α-helices and a small β-strands, with many residues (such as His^401^, His^403^, Tyr^427^, Glu^431^, Trp^438^, Arg^446^ and so on) on top, which might serve as potential substrate-binding sites. Finally, the C-terminal domain terminates at β17 in the second layer of the bilayer structure. Taken together, these structural elements constitute the C-terminal anti-parallel bilayer β-sheet domain of GAGase II (*Figure. 1A*).

Similar to GAGase II, GAGase VII also has an N-terminal (α/α)_6_ toroid domain (Leu^35^- Arg^360^) composed of 16 α-helices, but its α1 helix consists of only three residues (Glu^42^-Val^44^) and a long loop chain (*Figure. 1*A**). Moreover, the C-terminal two-layered β-sheet domain (Met^361^-Met^616^) of GAGase VII is also composed of 17 β-strands (β1-β15, β16a, β16b and β17). However, unlike GAGase II, its β16 strands is interrupted into two small β-strands (β16a, β16b) by a randomly coiled loop chain. Compared with GAGase II, these additional loop regions may make GAGase VII less structurally stable than GAGase II, and may be an important factor in why the activity of GAGase VII is far lower than GAGase II. Nevertheless, structural alignment further shows that GAGase II has good superposition with GAGase VII and other GAGases, indicating that GAGases are highly conserved in structures (*Supplemental Figure S1*).

### Mn binding site of GAGase II and GAGase VII

Unlike the GAG lyases identified from PL8, PL15 and PL21 families, which contain Ca^2+^ or Zn^2+^ in their overall crystal structures (***Shaya, et al., 2008***; ***Shaya, et al., 2006***; ***Zhang, et al., 2021***), a Mn^2+^ was detected in C-terminal domains of both GAGase II and VII by inductively coupled plasma-mass spectrometry (ICP-MS) analysis (*Figure 1B*, *Supplemental Table S1*). Structurally, the Mn^2+^ is located inside the long Ω loop after the β1-β5 antiparallel β-sheet, and coordinates with the nearby nitrogen and oxygen atoms of several residues such as Asp and His in both GAGase VII and GAGase II (GAGase II also has coordination with three surrounding water molecules), with distances of 2.0-2.5 Å. Such interaction should stabilize the nearby loop structures and may form a substrate-binding site relevant for substrate selectivity. To investigate the roles of these adjacent residues interacting with the Mn^2+^, His^403^, Asp^405^, Asp^421^ and His^457^ in GAGase II were individually mutated to alanine (Ala), and the results showed that these variants resulted in complete or substantial inactivation of GAGase II (*Figure 2A*). In addition, the inhibitory effects of the chelator (EDTA) and the stimulation of Mn^2+^ on the enzymatic activities also support the critical role of Mn^2+^ on GAGases (*Figure 2B*).

**Figure 2.**
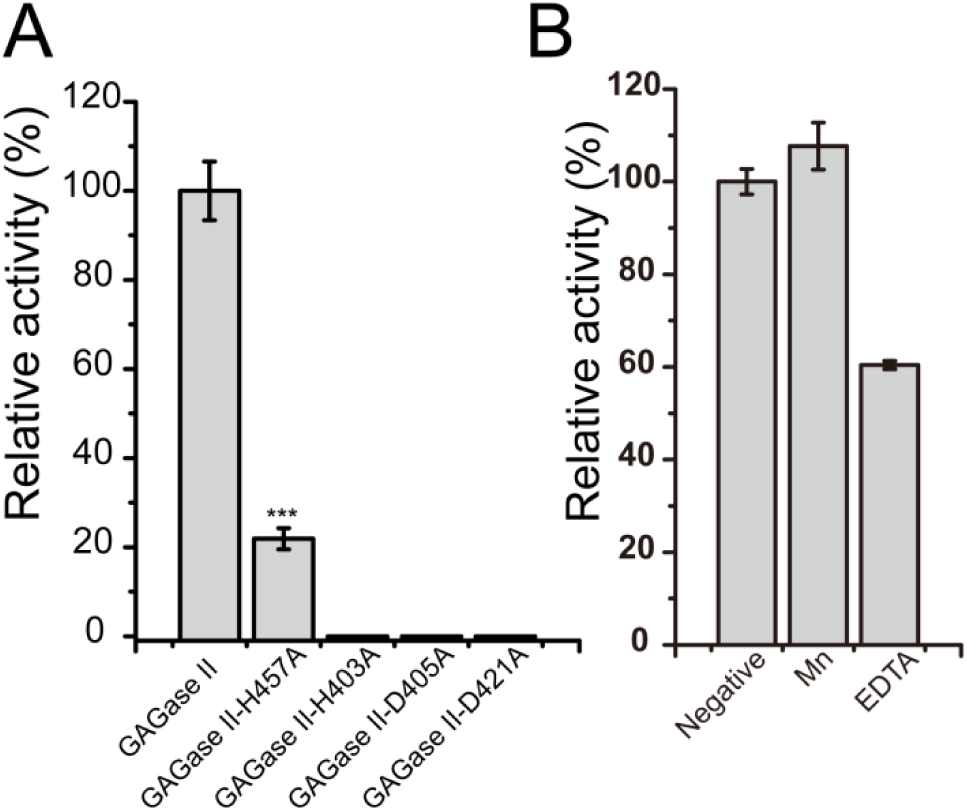
Activity of GAGase II and its variants. **(A)**, Activity of site-directed mutants of His and Asp nearby the Mn^2+^. His and Asp residues nearby the Mn^2+^ were individually mutated to Ala and the relative activity of each variant was measured as described in “*Materials and methods*”. All of the residual activities were evaluated and shown as the relative intensity compared with that of wild type GAGase II. ***: p<0.001. **(B)**, The effects of Mn^2+^ and chelating reagent of GAGase II were determined using HA (1 mg/ml) as substrate. Error bars represent averages of triplicates ± S.D.

### Multiple structural alignments of GAGase II and its structurally similar GAG lyases and alginate lyases

The ultrabroad substrate degradation spectra of GAGases indicate that they should be structurally similar with some other PLs with different substrate specificities. By using GAGase II as a representative, a structural alignment assay showed that the PL35 GAGases are structurally similar to various N-terminal (α/α)_n_ toroid domain and C-terminal antiparallel β- sheet domain-containing GAG/alginate lyases from PL8 (***Fethiere, et al., 1999***; ***Huang, et al., 2003***; ***Li and Jedrzejas, 2001***; ***Lunin, et al., 2004***; ***Shaya, et al., 2008***), PL12 (***Hashimoto, et al., 2014***; ***Ulaganathan, et al., 2017***), PL15 (***Ochiai, et al., 2010***; ***Zhang, et al., 2021***), PL17 (***Park, et al., 2014***), PL21 (***Shaya, et al., 2006***), PL23 (***Sugiura, et al., 2011***) and PL39 (***Ji, et al., 2019***). Notably, GAGase II shares very low sequence identity with PL21 heparinase II from *Pedobacter heparinus* (***Shaya, et al., 2006***) (23.1% identity with 29% query coverage), PL12 heparinase III from *Bacteroides thetaiotaomicronin* (***Ulaganathan, et al., 2017***) (23.6% identity with 32% query cover) and PL39 alginate lyase from *Defluviitalea phaphyphila* (***Ji, et al., 2019***) (26.5% identity with 31% coverage), and no significant sequence identity with PL17 alginate lyases (***Park, et al., 2014***), PL15 exoHep (***Zhang, et al., 2021***), PL8 chondroitin ABC lyase (***Shaya, et al., 2008***) and chondroitin AC lyase (***Lunin, et al., 2004***); however, all are relatively well superimposable in their overall structures and highly conserved in their triplet active site residues (*Figure 3*). In terms of overall structural similarities, the (α/α)_n_ toroid domain and anti-parallel β- sheet domain of GAGase II show more structural similarity to alginate lyases with (α/α)_6_ toroid and the antiparallel β-sheet domain, such as those from the PL15 (PDB code: 3A0O) (***Ochiai, et al., 2010***), PL17 (PDB code: 4NEI) (***Park, et al., 2014***) and PL39 (PDB code: 6JP4) (***Ji, et al., 2019***) families, than various GAG lyases (*Table 2,3*), indicating that GAGases might have originated from alginate lyases rather than GAG lyases. Reasonable support for this speculation could be the significant alginate-degrading activity of GAGase III. Therefore, the broad substrate spectrum of GAGases may be because they are evolutionarily transitional types with the enzymatic features of alginate lyases and GAG lyases.

**Figure 3.**
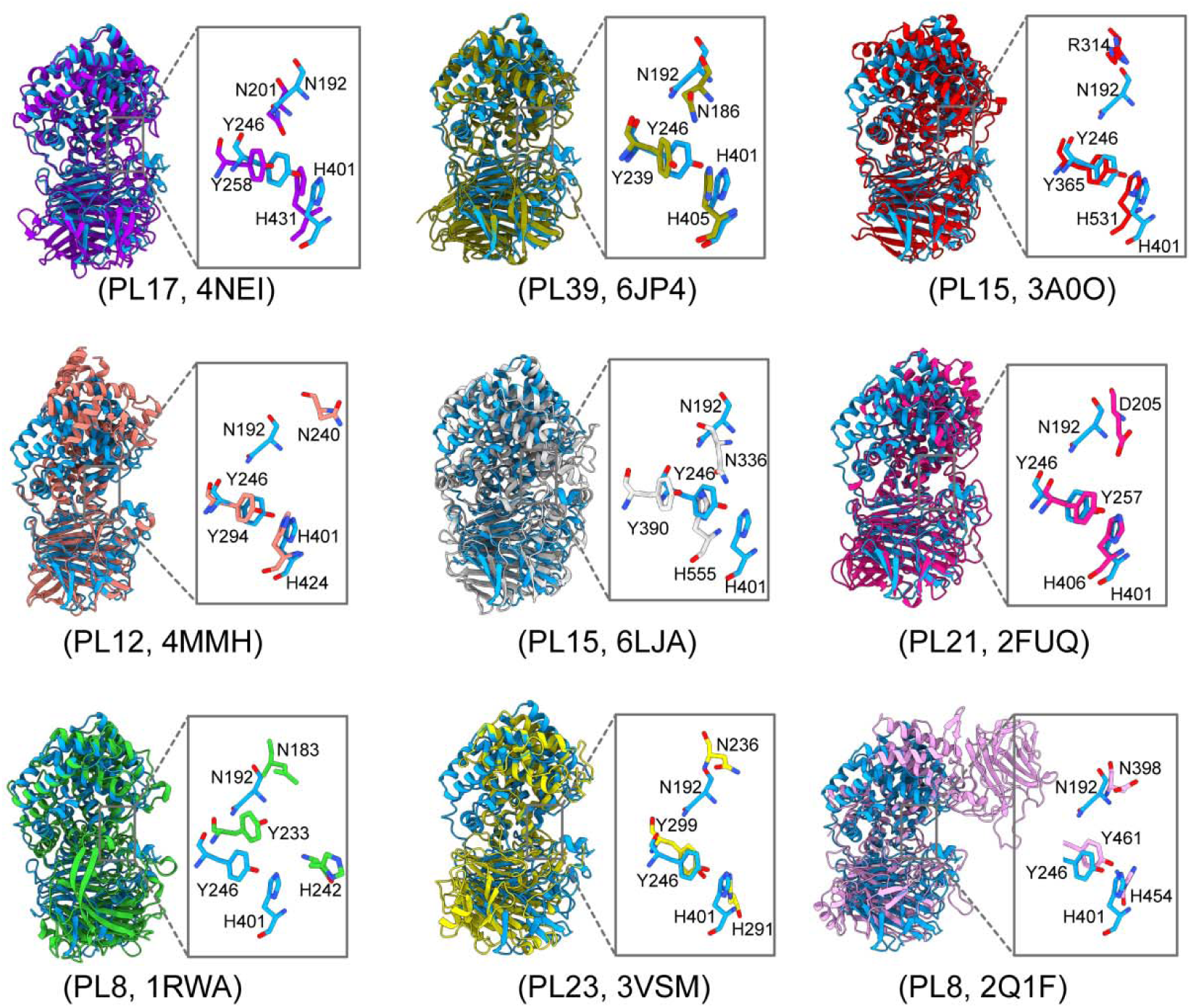
Multiple structural alignments of GAGase II and its structurally similar proteins. GAGase II (8KHV, *blue*) was aligned with structurally identified GAGs and alginate lyases, including PL17 family alginate lyase (4NEI, *purple*), PL39 family alginate lyase (6JP4, *olive*), PL15 family alginate lyase (3A0O, *red*), PL12 family heparinase III (4MMH, *salmon*), PL15 family exoHep (6LJA, *gray*), PL21 family heparinase II (2FUQ, *magenta*), PL8 family chondroitin sulfate AC lyase II (1RWA, *green*), PL23 family chondroitinase (3VSM, *yellow*) and PL8 family chondroitin sulfate ABC lyase II (2Q1F, *pink*).

**Table 2.**
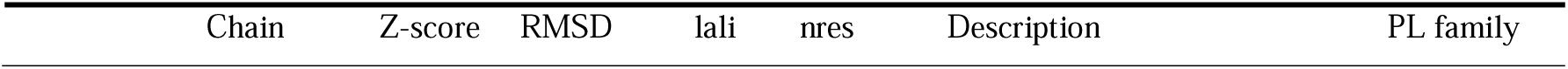
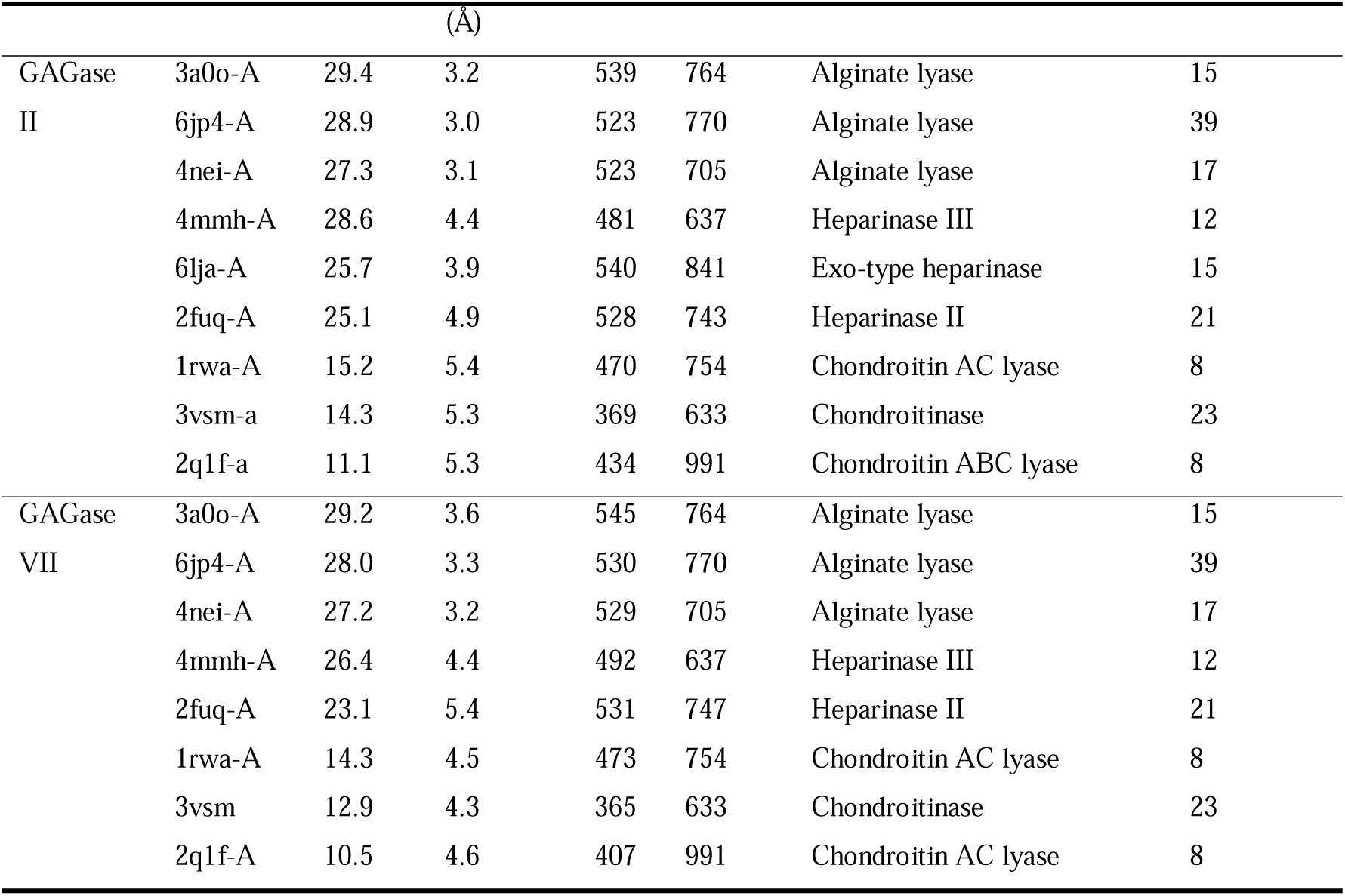
Structural similarity of GAGase II/GAGase VII with GAG/alginate lyases analysed using DALI.

**Table 3.**
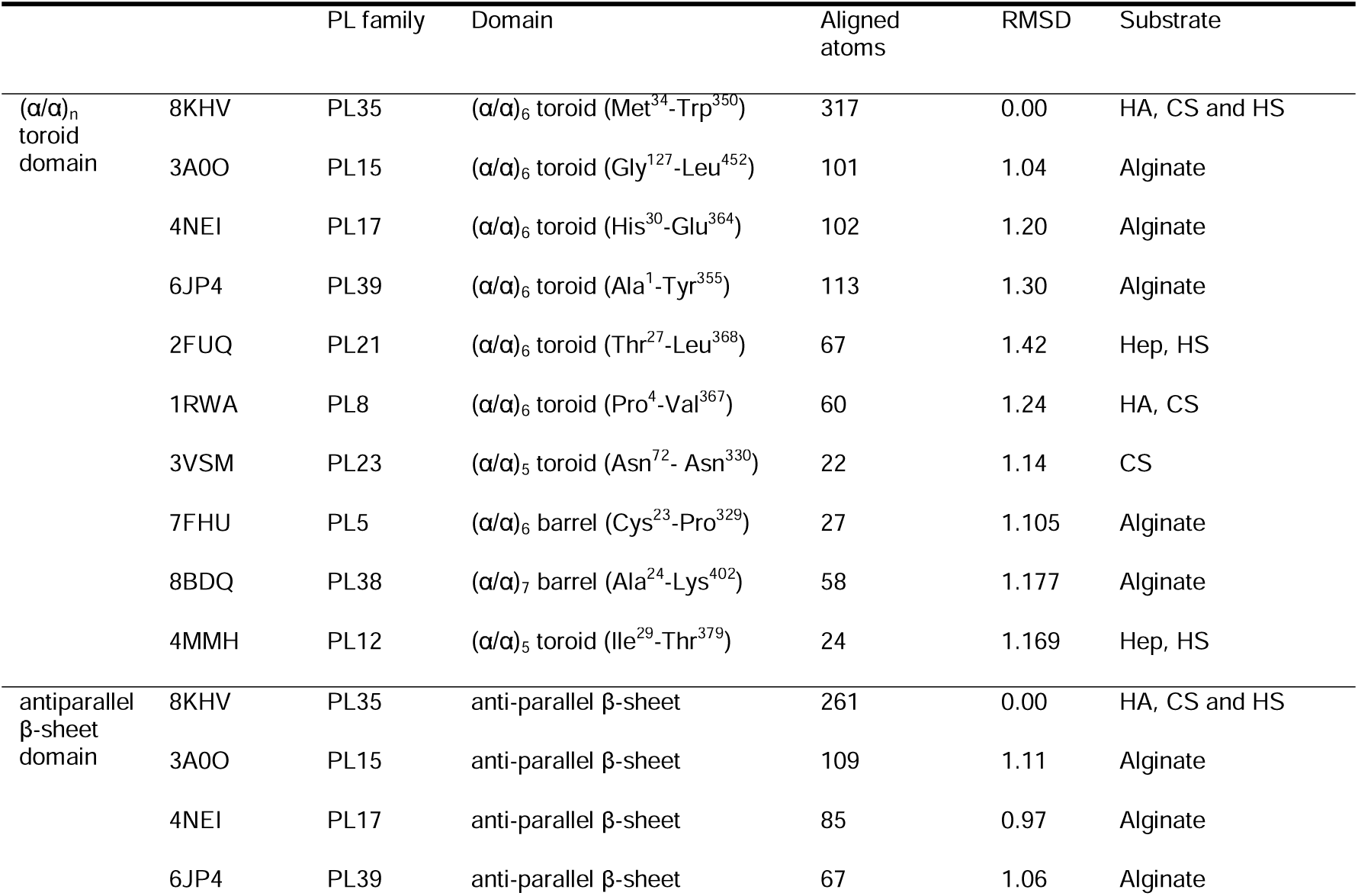
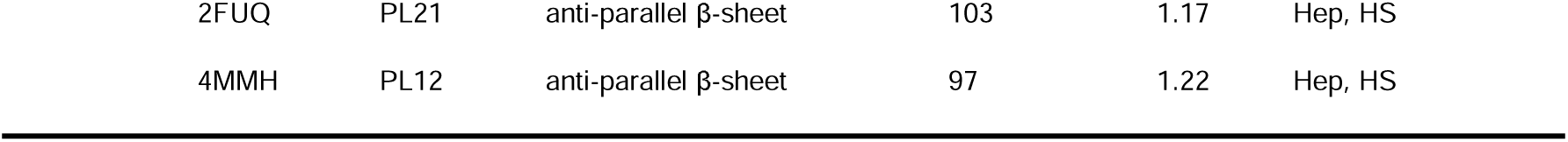
Multiple structural alignments of (α/α)_n_ toroid domain or antiparallel β-sheet domain of GAGase II and identified GAG/alginate lyases.

The detailed views of the crucial active site residues are shown in stick mode. The root-mean-square deviation (RMSD) between these structures and GAGase II were calculated as 1.34, 1.34, 1.30, 1.24, 1.15, 1.35, 1.24, 1.14 and 1.41 Å based on 171, 148, 141, 111, 99, 87, 60, 22 and 21 pruned atoms, respectively.

### Catalytic center and substrate binding sites of GAGases

Based on the sequence and structural alignment with homogenous alginate/GAG lyases, the conserved residues Tyr^246^, His^401^, Asn^192^ of GAGase II, Tyr^243^, His^398^, Asn^189^ of GAGase III, and Tyr^241^, His^397^, Asn^187^ of GAGase VII may be the key triplet residues involved in β-elimination catalysis (*Figure 3*). To confirm this hypothesis, these residues were individually replaced by Ala and the activity of the mutants was analyzed. The results showed that the activity of all variants toward CS-C was completely lost (*Figure 4A*), indicating that GAGases act through a general acid base catalytic mechanism by using His/Tyr as a Brønsted acid/base, similar to that observed for various other identified GAG/alginate lyases (*Table 4*). Compared with structurally similar GAG/alginate lyases, the results from molecular docking and catalytic tunnel predictions showed that GAGase II contains a shorter catalytic cavity (*Supplemental Figure S2*), which may explain why they can accommodate and degrade a variety of substrates with very different structures. However, we were unsuccessful in preparing the cocrystals of GAGases/their inactive variants with substrates to reveal their substrate binding and degradation mechanisms. Alternatively, molecular docking experiments of GAGase II with several structurally defined hexasaccharide (Hexa) substrates were performed as shown in *figure 4B*. Based on the docking results obtained for GAGase II with an HA Hexa (GlcUA1–3GlcNAc)_3_ (PDB code: 1HYA) (***Winter, et al., 1975***) (*Figure 4B*), the nonreducing end of the substrate at the “-1” and “-2” subsites is near some positively charged amino acids, including Arg^87^, Arg^88^, Arg^94^, His^136^, Asn^191^ and Asn^192^, in the vicinity of the catalytic cavity of GAGase II, which could facilitate the binding of negatively charged substrates. Besides, several acidic amino acids (Glu^242^, Asp^299^ and Glu^431^) are very close to the “+1” and “+2” subsites, which could promote the release of the reducing end product via charge repulsion. There are some other residues surrounding around the catalytic cavity, such as Tyr^240^, Gln^307^, Arg^342^, Tyr^427^, Glu^428^, Glu^431^, Trp^438^, Met^440^ and Arg^446^, which may play a key role in stabilization of the catalytic cavity and overall structure. Site-directed mutagenesis of these residues individually replaced with Ala showed that resulting variants were partially or completely inactivated (*Figure 4C*), confirming that these residues play important roles in the binding and release of substrate and structural stabilization.

**Figure 4.**
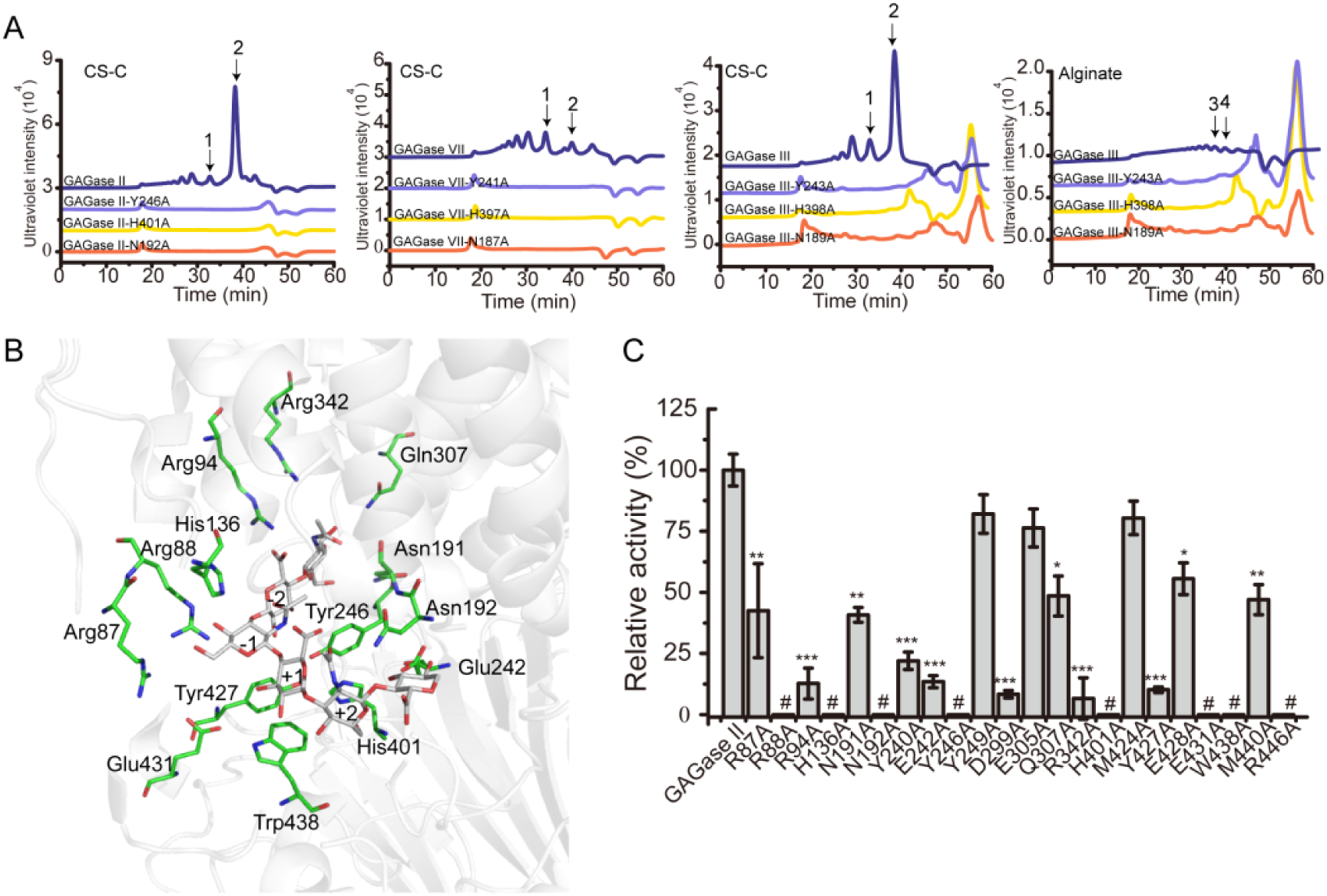
Catalytic center and substrate binding sites of GAGases. **(A)**, The crucial catalytic site-directed mutagenesis of GAGase II, GAGase III and GAGase VII. The conserved crucial residues of GAGase II, GAGase III and GAGase VII were individually mutated to Ala. CS-C and alginate were used as substrates for the activity evaluation of GAGase II, GAGase III, GAGase VII and its variants. The activity of each variant was detected using gel filtration HPLC on a Superdex Peptide column as described in “*Materials and methods*”; the elution of each fraction is indicated as follows: 1, CS-C tetrasaccharide; 2, CS-C disaccharide; 3, alginate trisaccharide; 4, alginate disaccharide. **(B)**, Molecular docking of GAGase II with a HA hexasaccharide. The molecule docking was carried out with GAGase II and a HA hexasaccharide (PDB code: 1HYA) to predict the substrate binding sites. The binding site residues (*green*) and hexasaccharide ligand (*gray*) are showed as sticks. **(C)**, The putative substrate-binding site residues surrounding the docking substrate were individually mutated to Ala. HA was treated with each variant at 40 °C for 12 h and the relative activity of each variant was shown as the relative intensity compared with that of wild type GAGase II. *: p<0.01, **: p<0.001, ***: p<0.0001, #: the activities of the variants were too low to be accurately detected and almost completely inactivated.

**Table 4.**
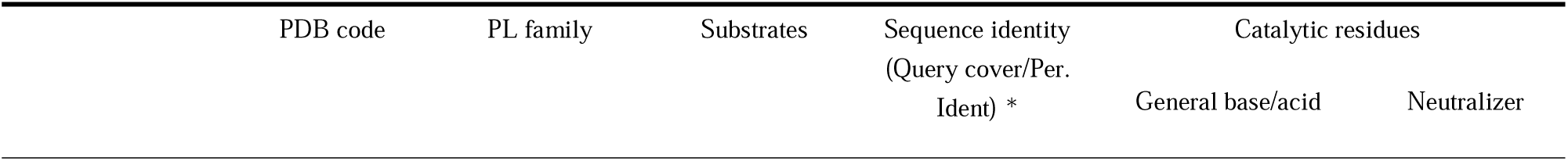
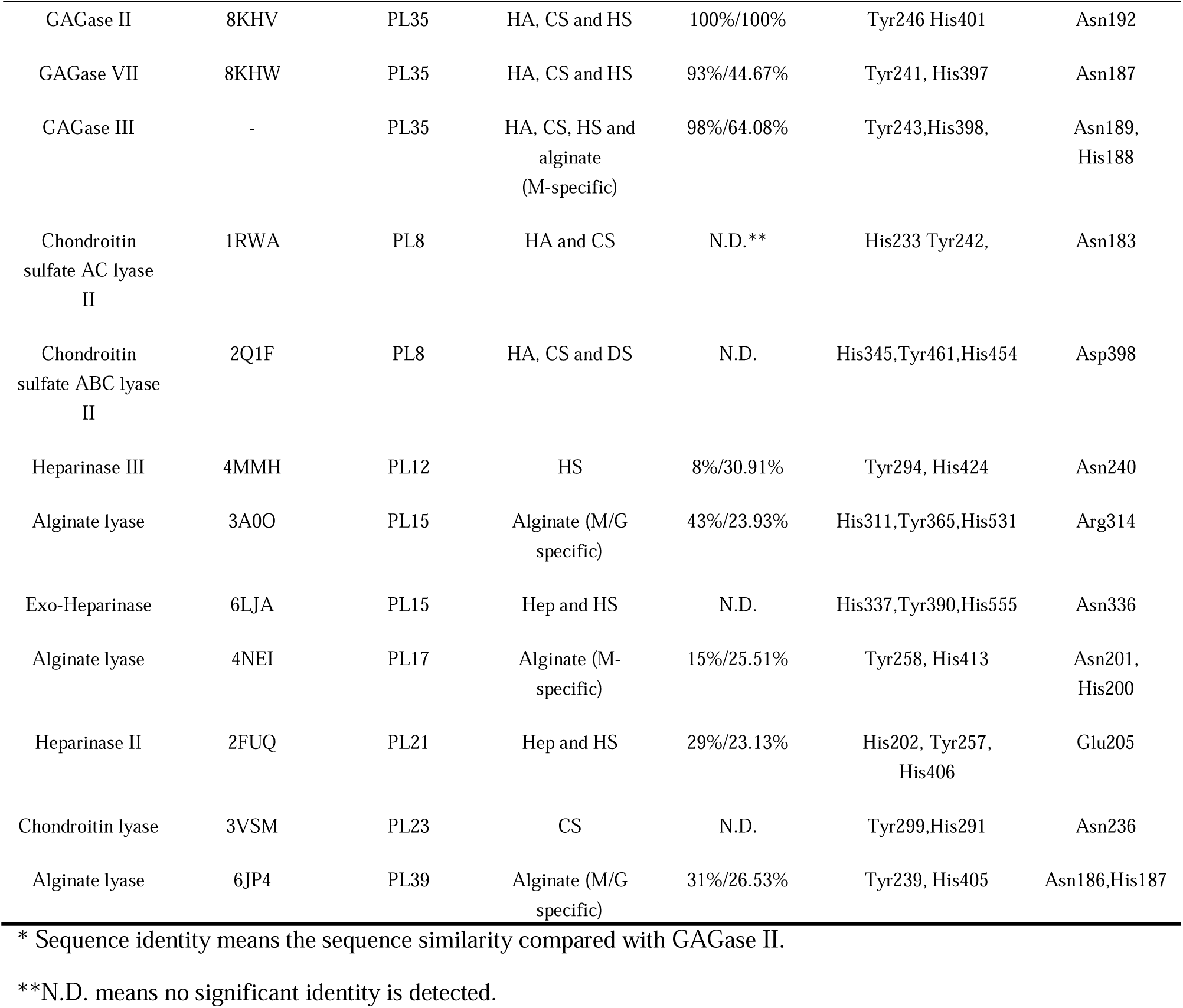
Catalytic residues comparison of identified GAGs and alginate lyases shared similar structure with GAGase II and GAGase VII.

### Substrate selectivity of GAGases

As mentioned above, GAGase III can degrade not only HA, CS and HS but also alginate, even though the activity on alginate is relatively low (about 1/50 of the activity on HA). To investigate the reason why this enzyme is capable to act on alginate, its structure was predicted using RoseTTAfold, and aligned with GAGase II and GAGase VII. Results showed that GAGase III has a unique His^188^ residue in its catalytic cavity, which is conserved in alginate lyases from PL17 (***Park, et al., 2014***), PL39 (***Ji, et al., 2019***), PL38 (***Ronne, et al., 2023***), PL5 (***Pandey, et al., 2021***) and PL15 (***Ochiai, et al., 2010***) families, while other GAGases have a conserved asparagine residue such as the Asn^191^ of GAGase II and the Asn^186^ of GAGase VII in the corresponding position (*Figure 5A-B*). To verify the function of this histidine, His^188^ was replaced with alanine and asparagine, respectively. The results of substrate-specificity analysis showed that the alginate-degrading activity of both GAGase III-H188A and GAGase III-H188N were abolished even at a quite high ratio of the mutated enzyme to substrate such as 30 μg enzyme to 30 μg substrate (*Supplemental Figure S3A*), while their GAG-degrading activity was only partially affected, indicating that the His^188^ residue is essential for the alginate-degrading activity of GAGase III (*Figure 5C-D*). However, when the asparagine residue at the corresponding position of other GAGases was mutated to histidine, the obtained variants did not exhibit any alginate-degrading activity (*Supplemental Figure S3B*), suggesting that degrading activity of GAGase II remains to be determined outside of the His^188^ residue.

**Figure 5.**
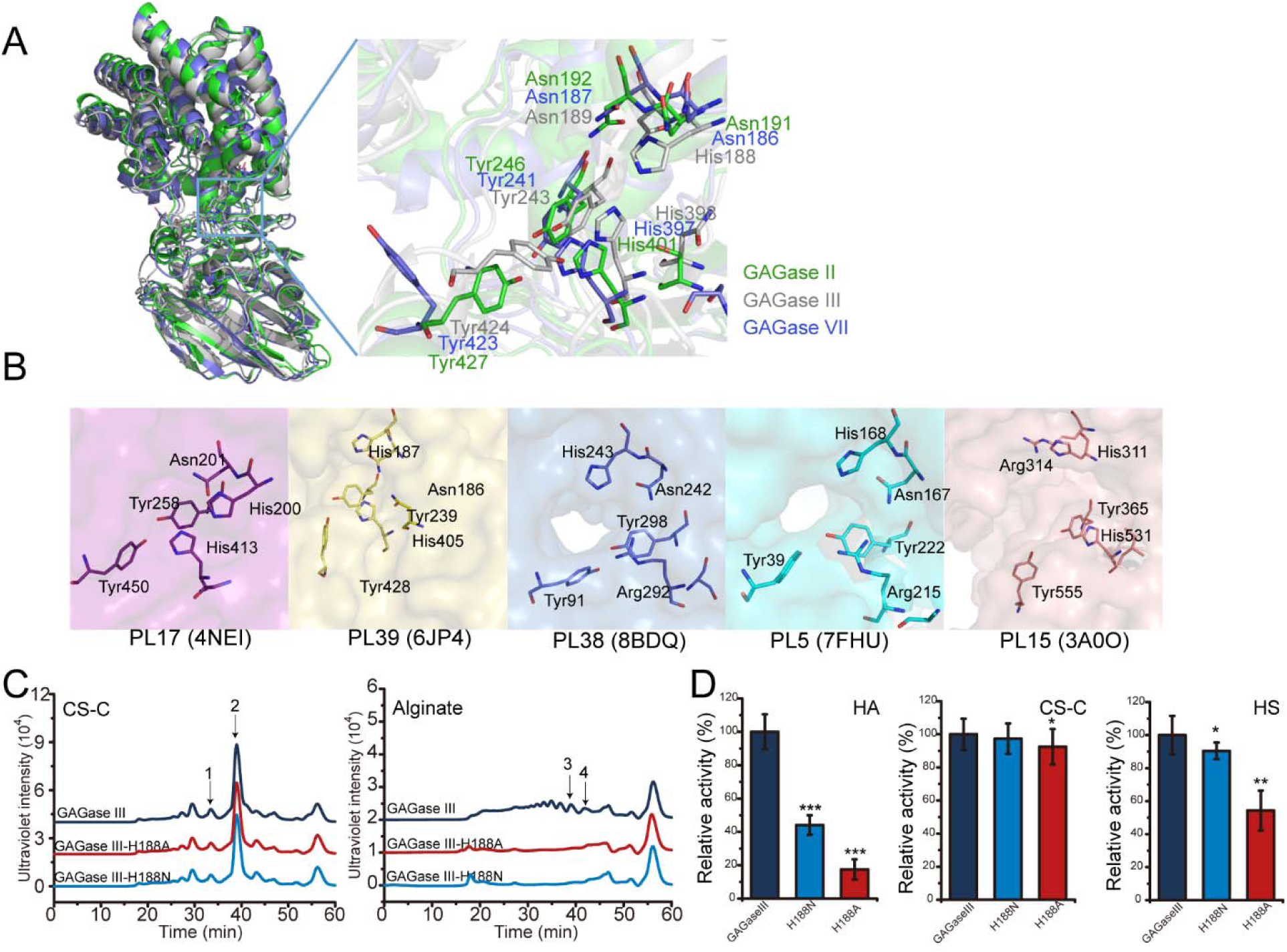
Analysis of a key residue for the alginate-degrading activity of GAGase III. (A), Multiple structural alignment of GAGase II, III and VII. GAGase II (8KHV, *green*) and GAGase VII (8KHW, *skyblue*) were aligned with GAGase III (model, *gray*). The detailed views of crucial catalytic residues are shown in stick mode; (B), Conserved catalytic residues of alginate lyases from PL17 (4NEI, *purple*), PL39 (6JP4, *yellow*), PL38 (8BDQ, *slate*), PL5 (7FHU, *cyan*) and PL15 (3A0O, *pink*) family. Residues are shown in stick; (C), Activity assay of GAGase III-H188N and GAGase III-H188A toward CS-C and alginate. The crucial site His^188^ was mutated to alanine and asparagine, respectively. The activity of each variant against CS-C and alginate was assessed using gel filtration HPLC on a Superdex Peptide column, as described under “*Materials and methods*”; the elution of each fraction is indicated as follows: 1, CS-C tetrasaccharide; 2, CS-C disaccharide; 3, alginate trisaccharide; 4, alginate disaccharide; (D), Relative activity of GAGase III and its variants. Three types of GAGs (HA, CS-C and HS) were individually treated with GAGase III and its variants (GAGase III-H188N and GAGase III-H188A) at 40 °C for 1 h; relative activities of enzymes were determined by detecting the absorbance at 232 nm. Data are shown as the percentage of the activity relative to the wild-type GAGase III. *, p<0.5; **, p<0.01; ***, p<0.001.

GAGases show preference for some specific sulfation patterns (***Wei, et al., 2024***). For example, GAGases prefer to degrade CS domains composed of non-/6-*O*-sulfated but not 4-*O*- sulfated GalNAc residues and D-GlcA residues. To further investigate the reason why these enzymes have such structural preference for substrates, we tried to prepare the co-crystal of GAGase II or VII with various structure-defined oligosaccharide substrates, but ultimately failed. Therefore, we used molecular docking analysis to preliminarily explore the possible reasons why GAGases have substrate structural preference. To this end, docking of GAGase II with a nonsulfated CS Hexa (GlcUAβ1–3GalNAc)_3_ (PDB code: 2KQO) and a tri-4-*O*-sulfated CS Hexa (GlcUAβ1–3GalNAc(4S))_3_ (PDB code: 1C4S) (***Sattelle, et al., 2010***) were carried out, respectively. In the docked model, the monosaccharide residues of the substrate were named “+1” and “+2” toward reducing end, and “-1” and “-2” toward non-reducing end from the β1-4 cleavage site following the nomenclature (***Davies, et al., 1997***). The results show that for the nonsulfated CS Hexa, the GlcUA residue at the “+1” subsite is closely approached by the Tyr^246^ residue as a Brønsted base (*Figure 6 left*), which facilitates the uptake of the C5 proton of GlcUA by Tyr^246^, and in the case of tri-4-*O*-sulfated CS Hexa, the 4-*O*-sulfate group in GalNAc residue at the “+2” subsite interacts with the neutralizer Asn^192^ and the catalytic residue Tyr^246^ via hydrogen bonds etc., the C5 proton of GlcUA at the “+1” subsite is away from the key catalytic residue Tyr^246^ (*Figure 6 right*, *Supplemental Figure S4*), which hinders the degradation of CS with such sulfation pattern, as we detected in the biochemical analysis above. Of course, the results from docking assay need to be further confirmed by more structural and biochemical evidence.

**Figure 6.**
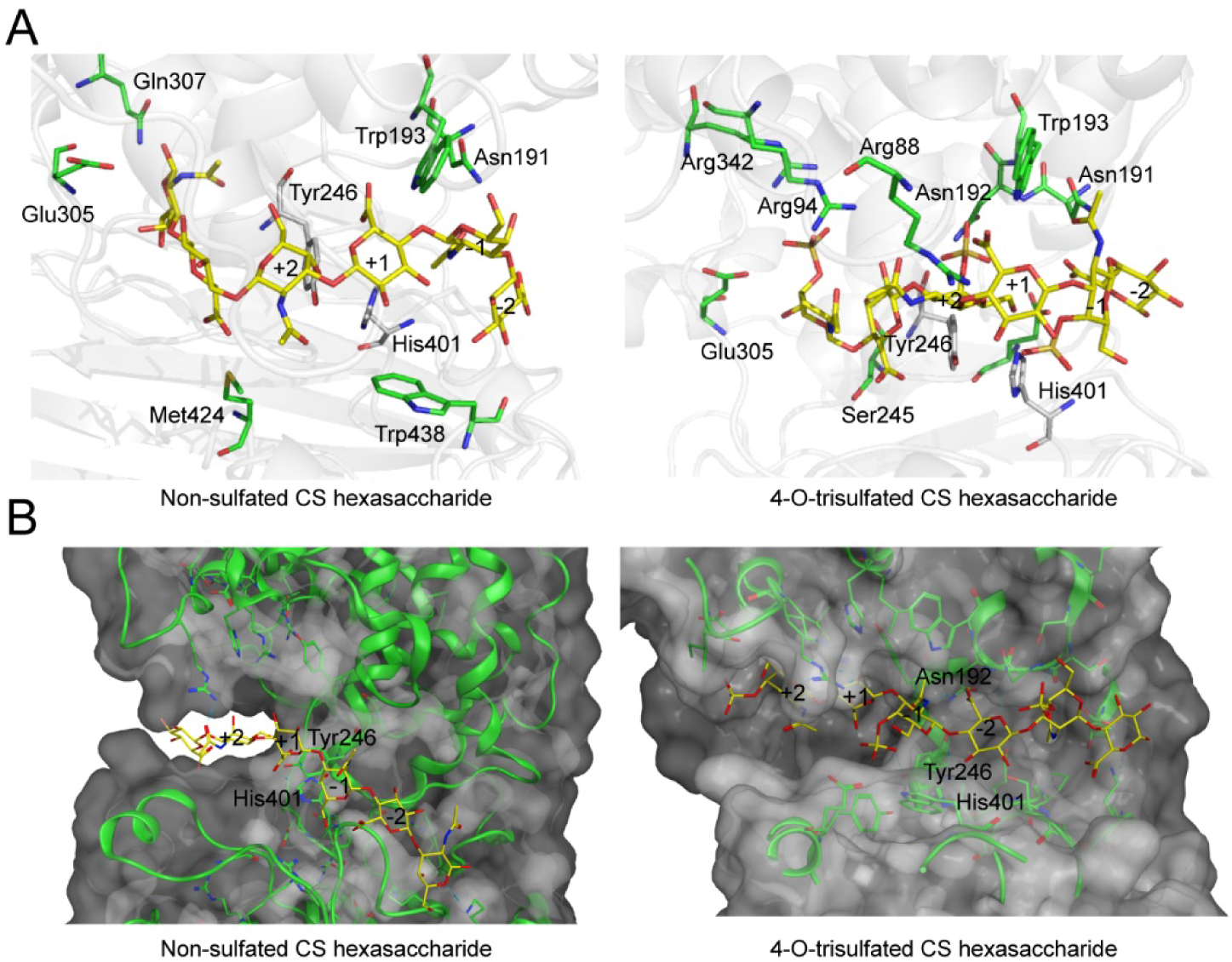
Molecular docking of GAGase II with hexasaccharide ligands. Molecular docking of GAGase II with CS ligands. The molecule docking was carried out with GAGase II and CS ligands, including nonsulfated (PDB code: 2KQO) (*left*) and 4-*O*-trisulfated (PDB code: 1C4S) (*right*) CS hexasaccharide, to investigate the substrate selectivity, using AutoDock Vina (A) and Molecular Operating Envioronment (MOE) (B). The catalytic triplet residues (*green*) and CS ligand (*yellow*) are showed as sticks.

### Catalytic mechanism of GAGases

In summary, taking GAGase II as representative, the catalytic process of GAGases is proposed as follows. The negatively charged GAG substrate binds to several positively charged residues, such as Arg^87^, Arg^88^, Arg^94^, His^136^, Asn^191^ and Asn^192^, near the catalytic triad residues in the catalytic cavity. The Asn^191^ and Asn^192^ residues interact with the carboxyl group of GlcUA at the +1 subsite to reduce the p*K*a and promote the protonation of the C5 proton. The His^401^ works as a proton receptor (Brønsted base) to abstract protons from the C5 position of GlcUA and create an enolate tautomeric. The protonated Tyr^246^ (Brønsted acid) provides a proton to the C-4–O-1 glycosidic bond between the “-1” and “+1” subsite. The glycosidic bond is cleaved as an electron transfer occurs from the carboxyl group to form a double bond between C4 and C5 of GlcUA. A new reducing end and a new unsaturated nonreducing end are recreated. Moreover, the degradation of polyM by GAGase III requires the involvement of His^188^ residue, which may involve in the neutralization of carboxyl groups in M-blocks in alginate (*Figure 7*).

**Figure 7.**
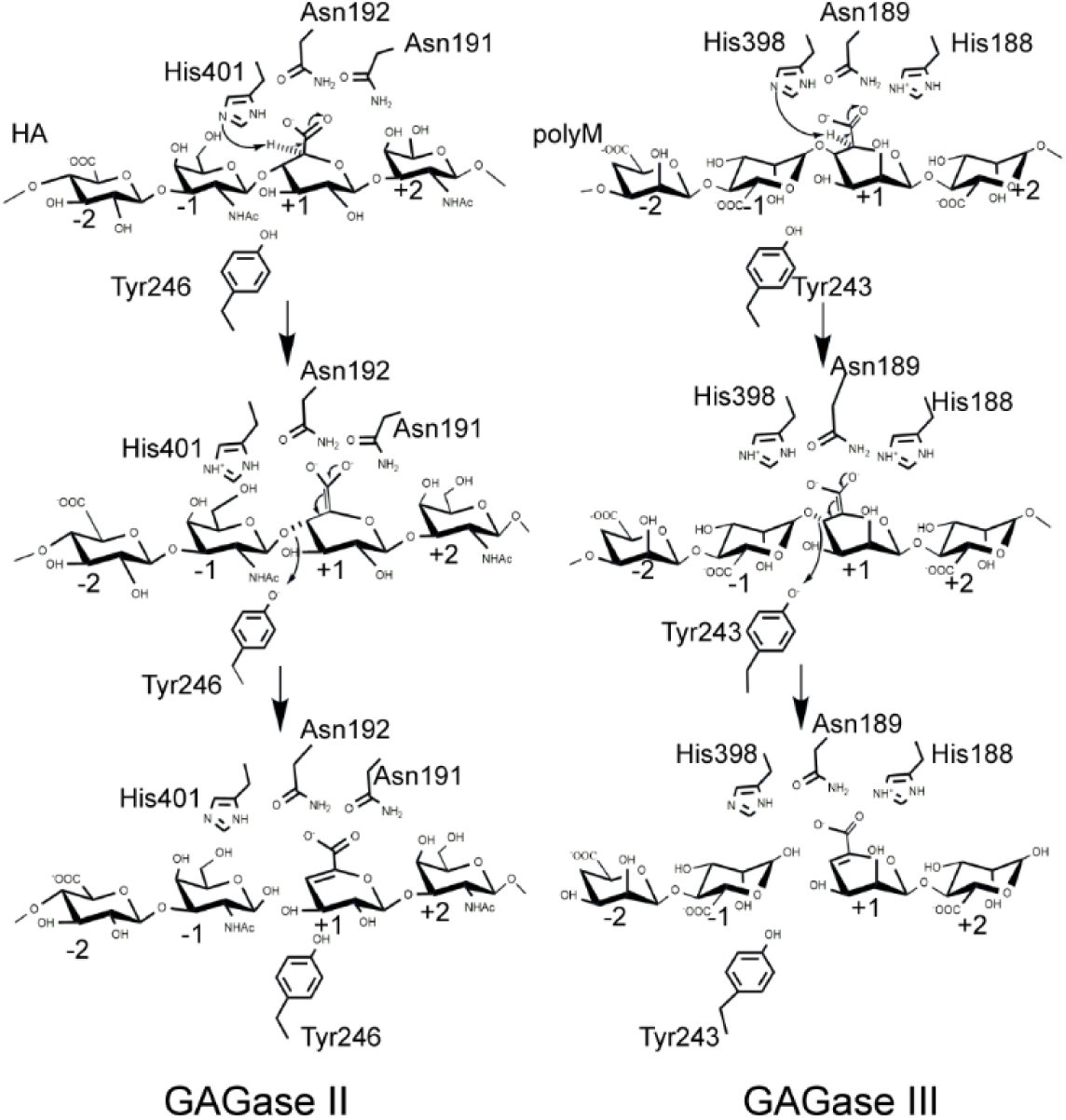
Proposed catalytic mechanism of GAGase II and GAGase III. Take the HA degradation by GAGase II and the polyM degradation by GAGase III for example. Briefly, the substrate is firstly binding to the negatively charged residues near the catalytic sites, the carboxylate group is neutralized by Asn^192^, Asn^191^ in GAGase II and His^188^, Asn^189^ in GAGase III; His^398^ in GAGase II and His^401^ in GAGase III are proposed to work as general base to abstract a proton from the C5 position of GlcUA at +1 position, and Tyr^246^ in GAGase II and Tyr^243^ in GAGase III work as general acid to donate the leaving group at -1 position a proton. The arrows indicate the direction of electron transfer, and the C5-C6 double bond on the middle panel indicate the enolate anion intermediate created by proton abstraction at C-5 position.

## Discussion

GAG lyases derived from microorganisms, as GAG-specific degrading enzymes, are classified into 13 PL families in the CAZy database (***Drula, et al., 2022***). They are not only critical for microorganisms to degrade and utilize GAGs as carbon sources, but also become useful tools for structural and functional studies of GAGs. In contrast, alginate lyases, as alginate-specific degrading enzymes, are classified into 15 PL families in the CAZy database (***Drula, et al., 2022***). They are essential for the metabolism of alginate in microorganisms and algae as well as lower marine animals, and have completely different substrate specificities than GAG lyases. Due to the similarity between GAG lyases and alginate lyases in 3D fold, GAG lyases are thought to have originated from the divergent evolution of alginate lyases (***Garron and Cygler, 2014***), but enzymes that exhibit transitional features in function and structure, which would provide direct evidence to support this view, are lacking. On the basis of previous functional studies (***Wei, et al., 2024***), structural studies of GAGases with HA, CS, HS and even alginate-degrading activities should further provide substantial evidence for the evolution of GAG lyases.

Prior to this study, no member of the PL35 family had been structurally characterized. Here, the structures of GAGase II and VII were determined at resolutions of 1.9 Å and 2.4 Å, respectively, and both exhibit a highly similar structure of “N-terminal (α/α)_6_ toroid and C-terminal two-layer antiparallel β-sheet”, which is also the fold adopted by most structurally known GAG lyases except CSase B in PL6 (***Huang, et al., 1999***), Hepase I in PL13 (***Han, et al., 2009***) and hyaluronan lyase in PL16 (***Martinez-Fleites, et al., 2009***). Structural alignment has shown that GAGases share considerable degrees of similarity with all GAG/alginate lyases with the (α/α)_n_ toroid and antiparallel β-sheet fold in PL8 (***Fethiere, et al., 1999***), PL12 (***Hashimoto, et al., 2014***), PL15 (***Ochiai, et al., 2010***; ***Zhang, et al., 2021***), PL17 (***Park, et al., 2014***), PL21 (***Shaya, et al., 2006***), PL23 (***Sugiura, et al., 2011***) and PL39 (***Ji, et al., 2019***) families, especially the key catalytic triplet residues (Tyr, His and Asn) in their active centers, which are highly conserved; thus, they may share a similar Brønsted base/acid catalytic mechanism (***Garron and Cygler, 2010***; ***Garron and Cygler, 2014***). Notably, GAGases are more structurally similar to alginate lyases from PL15 (***Ochiai, et al., 2010***), PL17 (***Park, et al., 2014***) and PL39 (***Ji, et al., 2019***) family than to various GAG lyases, which further structurally supports the speculation that GAGases originated from alginate lyases.

As mentioned previously, many different GAG/alginate lyases share similar catalytic sites and the same catalytic machinery (***Garron and Cygler, 2010***; ***Garron and Cygler, 2014***), but their differences in substrate selectivity and endo-/exolytic manner are mainly attributed to subtle differences in the three-dimensional structure (***Lombard, et al., 2010***), which may be the substrate-binding sites. The recognition of appropriate substrates by PLs mainly depends on the interaction between the positively charged side chains of basic amino acid residues in their substrate-binding sites and the negatively charged carboxyl groups of HexUA residues in substrates. Moreover, compared to the structurally and biochemically characterized members of the PL8, PL12, PL15, PL21 and PL39 families, PL35 family proteins possess a shorter catalytic cavity; thus, PL35 family proteins may more easily accommodate various substrates with different structures but may exhibit a weaker ability to bind and degrade specific substrates, resulting in the activities of GAGases towards various GAGs and alginate are much lower than those of GAG or alginate-specific lyases. Certainly, the accurate substrate selectivity mechanism of GAGases remains to be revealed by preparing and resolving co-crystals of the enzyme-substrate complex in the future.

In addition, GAGases tend to act on GlcUA-containing HA, CS and HS but not IdoUA- containing DS and Hep, which should result from the absence of a His residue functioning as an *anti*-base in the catalytic cavity (***Garron and Cygler, 2010***; ***Shaya, et al., 2008***; ***Shaya, et al., 2006***). The His residue as an *anti*-base can substitute Tyr to absorb the C5 proton in the IdoUA and ensure that β-elimination is carried out successfully, which is detected in chondroitin sulfate ABC lyase (2Q1F) from PL8 family (His^454^) (***Shaya, et al., 2008***) and heparinase II (2FUQ) from PL21 family (His^202^) (***Shaya, et al., 2006***), and endows these enzymes with a bifunctional activity against GlcUA- and IdoUA-containing GAGs. Notably, GAGase III, the only identified GAGase with alginate-degrading activity, has a unique His^188^ residue also conserved in various alginate lyases from PL5, PL15, PL17, PL38 and PL39 families. Mutation assay showed that the His^188^ is a key residue for the alginate-degrading activity of GAGase III. However, when a conversed asparagine residue at the corresponding position of other GAGases was replaced by histidine, the muted GAGases still could not act on alginate, indicating that some other unidentified structural factors are also involved in the alginate degradation of GAGase III. Overall, while the substrate-degrading features of GAGases can be preliminarily elucidated structurally, more direct evidence remains to be obtained by detailed structural and biochemical approaches.

In addition, the identification of GAGases may help clarify an inappropriate naming issue regarding the functional module “Hepar_II_III superfamily”. This functional module is commonly found in not only GAG lyases but also alginate lyases with an “N-terminal (α/α)_n_ toroid + C-terminal β-sandwich” structure. Interestingly, however, a large number of identified alginate lyases with the functional module “Hepar_II_III superfamily” do not have the ability to degrade Hep/HS. Historically, due to the importance of Hep as an anticoagulant in clinical application, Hep/HS-related studies including heparinases have received much higher attention than alginate-related studies including alginate lyases, which may be the reason why some sequences in alginate lyases that are similar to heparinase are annotated named the “Hepar_II_III superfamily” module. From an evolutionary perspective, this “Hepar_II_III superfamily” module might be a conserved functional domain in the divergent evolution process from alginate lyase to GAG lyase. Therefore, it might be more reasonable and accurate for this functional domain to be named as the “alginate lyase superfamily” module, and the presence of this functional module in GAGases with evolutionary transitional characteristics may well support this view.

In conclusion, the first determination of the structure of GAGases not only facilitates the elucidation of the catalytic mechanism of PL35 family enzymes, especially the substrate selection mechanism of GAGases, but also provides potential evidence for the hypothesis that GAG lyase evolved from alginate lyase.

## Materials and methods

### Materials

HA from *Streptococcus equi*, CS-C from shark cartilage, alginate sodium from brown algae, polyethylene glycol (PEG) 6000, PEG 400, calcium chloride (CaCl_2_), tris-(hydroxymethyl) aminomethane (Tris) and hydroxyethyl piperazine ethanesulfonic acid (HEPES) were purchased from Sigma Aldrich. Lysine, phenylalanine, threonine, leucine, isoleucine, valine and L-selenomethionine were purchased from Solarbio Co.,Ltd. (Beijing, China).

### Heterologous expression and purification of GAGases and GAGase II selenomethionine derivant

The cloning and induced expression of PL35 family GAGases were described in previous study (***Wei, et al., 2024***). Briefly, the synthesized expression plasmid pET-30a-GAGases were transformed into *E. coli* BL21 (DE3) cells and culture in LB broth at 37 °C. Recombinant enzymes with a 6His-tag were expressed at 16 °C for 16 h by inducing with isopropyl 1-thio-β- D-galactopyranosid (IPTG) (final concentration of 0.05 mM) at the cell density of OD_600_=0.6–0.8. Furthermore, to obtain the Se-GAGase II, cultured cells harboring the pET-30a-GAGase II were centrifuged at 3,700 g at 4 °C for 5 min, washed with M9 culture medium (17.2 g/L Na_2_HPO_4_, 3 g/L KH_2_PO_4_, 2.5 g/L NaCl and 5 g/L NH_4_Cl), transferred into M9 culture medium supplemented with 1 mM MgSO_4_, 0.1 mM CaCl_2_, 0.4% (w/v) glucose and cultured at 37 °C until the OD_600_ reached 0.6. Then, the culture medium was cooled to 22 °C. Additional essential amino acids, including lysine (100 mg/L), phenylalanine (100 mg/L), threonine (100 mg/L), leucine (50 mg/L), isoleucine (50 mg/L), valine (50 mg/L), and L-selenomethionine (L-SeMet) (50 mg/L), were added and incubated at 22 °C for 15 min. The proteins contained L-SeMet were also induced and expressed at 16 °C for 16 h by supplementing with 5 mM IPTG.

After further induced cultivation, cells were harvested, resuspended and disrupted by sonication, the recombinant proteins in the supernatant of cell lysates are primarily purified by loading on a nickel affinity column, washing with buffer A containing 10 mM imidazole to remove impurities, and finally eluting using buffer A containing 250 mM imidazole. After nickel affinity chromatography, the elutes of GAGase II and GAGase VII were diluted and loaded on a Q-Sepharose FF column (GE Healthcare) eluted with a gradient concentration of NaCl from 0 to 1 M, and the target proteins were desalted and concentrated through ultrafiltration and further sub-fractionated by gel filtration on a Superdex G-200 column (GE Healthcare) for crystal culture. The concentration of each protein was determined using BCA Protein Assay Kit with bovine serum albumin (BSA) as reference protein (Cwbio, Shanghai).

### Activity assay of GAGase II, III, VII and their variants toward various GAGs and alginate

To evaluate the activity of GAGase II, III, VII and their variants, CS-C or alginate (30 μg) was treated with each wild-type enzyme or its variant (6 μg) in the optimal buffer (50 mM Tris-HCl, pH 7.0) for 12 h at 40 °C. In order to evaluate the activity of GAGase III-H188N and GAGase III-H188A in degrading alginate, alginate (30 μg) was treated with each variant (3-30 μg) at 40 °C for 12 h. After inactivated by boiling for 10 min, the products were analyzed by gel filtration HPLC using a Superdex^TM^ Peptide 10/300 GL column eluted with 0.20 M NH_4_HCO_3_ at a flow rate of 0.4 ml/min and monitored at 232 nm using a UV detector, and online analysis was conducted using LCsolution version 1.25.

To evaluate the activity of variants from the site-directed mutagenesis of some potential substrate-binding residues of GAGase II, HA (150 μg) was treated with the wild-type or each mutant (6 μg) in the optimal buffer (50 mM Tris-HCl, pH 7.0) at 40 °C for 12 h. To evaluate the activity of GAGase III-H188N and GAGase III-H188A, HA, CS-C, or HS (150 μg) was treated with GAGase III and each variant (6 μg) at 40 °C for 1 h. The absorbance of each resultant was determined at 232 nm using a UV spectrophotometer.

### Crystallization, X-ray diffraction and data collection of GAGase II, GAGase VII and Se- Met-GAGase II

The purified GAGase II (20 mg/ml) was crystallized in 0.1 M HEPES (pH 7.0), 12% (w/v) PEG 6000 and 0.2 M CaCl_2_. The purified Se-Met-GAGase II (5 mg/ml) was crystallized in 0.1 M Tris (pH 7.5), 20% (w/v) PEG 6000 and 0.2 M CaCl_2_. The purified GAGase VII (10 mg/ml) was crystallized in 0.1 M HEPES (pH 7.0) and 36% (v/v) PEG 400. The crystallization of all the above proteins was carried out at 18 °C using hanging drop vapor diffusion method in 24-well plates. Crystals were harvested using nylon loops and soaked in cryoprotectant consisting of 20% glycerol and 80% mother liquor. X-ray diffraction data were collected at Shanghai Synchrotron Radiation Facility (SSRF) BL18U1 beamline. Diffraction datasets were processed using XDS (***Kabsch, 2010***). Data collection statistics are summarized in *Table 1*.

### Structure determination and refinement

The structure of GAGase II was solved by selenium single-wavelength anomalous dispersion (Se-SAD) using AutoSol from Phenix suite (***Adams, et al., 2011***). The structure of GAGase VII was determined by molecular replacement using Phaser (***McCoy, et al., 2007***) with the structure of GAGase II as the search model. Initial model building was carried out using AutoBuild from Phenix suite. Then several rounds of refinement and manual building were performed alternately using Phenix.Refine and Coot (***Emsley and Cowtan, 2004***), respectively. The ion type was confirmed by inductively coupled plasma-mass spectrometry (ICP-MS) analysis. For each sample, 10 ml of GAGase II or GAGase VII (10 mg) was mixed with 10 ml of nitric acid in equal proportions. The mixture was then placed in a metal bath at 120 °C and heated until clarified. Subsequently, the clarified mixture was filtered through a 0.22 µm membrane and analyzed using ICP-MS (NexlON, PerkinElmer). The structures of other GAGases were predicted using RoseTTAfold (***Baek, et al., 2021***). Structure alignments were performed using ChimeraX with point accepted mutation (PAM)-120 mtrix (***Pettersen, et al., 2021***). All of the structure illustrations were generated using ChimeraX (https://www.cgl.ucsf.edu/chimerax/) and Pymol (https://www.pymol.org/2/). A structure-based similarity search for GAGase II and VII were performed using the DALI server (http://ekhidna2.biocenter.helsinki.fi/dali/) (***Holm and Rosenstrom, 2010***). The identification of tunnels and channels in proteins were performed using the Caver Web v1.2 (https://loschmidt.chemi.muni.cz/caverweb/) (***Stourac, et al., 2019***). The classification of identified enzymes was confirmed based on the database of carbohydrate-active enzymes (CAZy) (http://www.cazy.org/) (***Drula, et al., 2022***) and the structures of identified PLs were download from the RCSB Protein Data Bank (PDB) (https://www.rcsb.org/) (***Berman, et al., 2000***).

### Site-directed mutagenesis of GAGases

Some potentially important residues located in and around the active center were individually replaced with alanine (Ala) by using the Fast Mutagenesis Kit V2 from Vazyme Biotech Co., Ltd (Nanjing, China) and these mutants were expressed and examined for activity against various substrates (HA, CS-C, HS, and alginate) to confirm their crucial role in enzyme catalysis. The primer pairs of these mutants are shown in the *Table S2*. The expression and purification of GAGases and each variant were estimated by SDS-polyacrylamide gel electrophoresis (SDS-PAGE) followed by staining with Coomassie Brilliant Blue R-250. The results of SDS-PAGE analysis are shown in *Supplemental Figure S5*.

### Molecular docking

Tertiary structure-defined GAG or alginate hexasaccharide was used to dock into the active cavity of GAGase II by using the Autodock Vina program (***Eberhardt, et al., 2021***; ***Trott and Olson, 2010***) and Molecular Operating Envioronment. Docking by using the Autodock Vina program was performed on a search space of 15.00×18.19×19.53 Å covering the active centre (x, y, z = -2.60, -12.57, 34.13). The binding energies using non-sulfated and 4-*O*-trisulfated hexasaccharides are calculated as -5.874 and -5.163 kcal/mol, respectively.

## Data availability

All data supporting the findings of this study are available within the paper (and supporting information files). All relevant data generated during this study or analysed in this published article are available from the corresponding author on reasonable request. The atomic coordinates and structure factors of the structures in this study have been deposited in the Protein Data Bank (PDB codes: 8KHV and 8KHW). The sequences of GAGase II (GenBank: SOD82962.1) and GAGase VII (GenBank: EDV05210.1) have already existed in NCBI database.

## Acknowledgment

This work was supported by the National Key R&D Program of China (No. 2021YFC2103100), National Natural Science Foundation of China (Nos. 31971201, 32330001, 31570071, 31800665), Major Scientific and Technology Innovation Project (MSTIP) of Shandong Province (No. 2019JZZY010817), Natural Science Foundation of Shandong Province (No. ZR2023MC017), the SKLMT Frontiers and Challenges Project (SKLMTFCP-2023-06), Shandong Provincial Youth Innovation Science and Technology Support Program for Colleges and Universities (2022KJ003).

## Author contributions

Wei Lin, Investigation, Validation, Visualization, Writing-Original Draft; Cao Hai-Yan, Investigation, Validation, Visualization, Writing-Original Draft; Zou Ruyi, Investigation, Validation; Du Min, Investigation, Validation; Zhang Qingdong, Investigation; Lu Danrong, Resources; Xu Xiangyu, Investigation; Xu Yingying, Investigation; Wang Wenshuang, Investigation; Chen Xiu-Lan, Investigation; Zhang Yu-Zhong, Conceptualization, Methodology, Writing-Reviewing and Editing; Li Fuchuan, Conceptualization, Methodology, Writing-Reviewing and Editing.

## Notes

### Competing Interest Statement

The authors have declared no competing interest.

### Summary of Updates

This manuscript is a revised version

## References

1. Adams PD, Afonine PV, Bunkóczi G, Chen VB, Echols N, Headd JJ, Hung L, Jain S, Kapral GJ, Grosse Kunstleve RW, McCoy AJ, Moriarty NW, Oeffner RD, Read RJ, Richardson DC, Richardson JS, Terwilliger TC, Zwart PH. 2011. The Phenix software for automated determination of macromolecular structures. Methods 55:94–106. doi: 10.1016/j.ymeth.2011.07.005

2. Baek M, DiMaio F, Anishchenko I, Dauparas J, Ovchinnikov S, Lee GR, Wang J, Cong Q, Kinch LN, Schaeffer RD, Millan C, Park H, Adams C, Glassman CR, DeGiovanni A, Pereira JH, Rodrigues AV, van Dijk AA, Ebrecht AC, Opperman DJ, Sagmeister T, Buhlheller C, Pavkov-Keller T, Rathinaswamy MK, Dalwadi U, Yip CK, Burke JE, Garcia KC, Grishin NV, Adams PD, Read RJ, Baker D. 2021. Accurate prediction of protein structures and interactions using a three-track neural network. Science 373:871–876. doi:10.1126/science.abj8754

3. Berman HM, Westbrook J, Feng Z, Gilliland G, Bhat TN, Weissig H, Shindyalov IN, Bourne PE. 2000. The protein data bank. Nucleic Acids Res 28:235–242. doi:10.1093/nar/28.1.235

4. Charnock SJ, Brown IE, Turkenburg JP, Black GW, Davies GJ. 2002. Convergent evolution sheds light on the anti-beta-elimination mechanism common to family 1 and 10 polysaccharide lyases. P Natl Acad Sci Usa 99:12067–12072. doi:10.1073/pnas.182431199

5. Clementi F. 1997. Alginate production by *Azotobacter vinelandii*. Crit Rev Biotechnol 17:327–361. doi:10.3109/07388559709146618

6. Cohen E, Merzendorfer H. 2019. Extracellular sugar-based biopolymers matrices. Springer International Publishing AG, Cham doi:10.1007/978-3-030-12919-4

7. Csoka AB, Stern R. 2013. Hypotheses on the evolution of hyaluronan: A highly ironic acid. Glycobiology 23:398–411. doi:10.1093/glycob/cws218

8. Davies GJ, Wilson KS, Henrissat B. 1997. Nomenclature for sugar-binding subsites in glycosyl hydrolases. Biochem J 321 (**Pt 2**):557–559. doi:10.1042/bj3210557

9. Drula E, Garron M, Dogan S, Lombard V, Henrissat B, Terrapon N. 2022. The carbohydrate-active enzyme database: functions and literature. Nucleic Acids Res 50: D571–D577. doi:10.1093/nar/gkab1045

10. Dyer DP, Salanga CL, Volkman BF, Kawamura T, Handel TM. 2015. The dependence of chemokine– glycosaminoglycan interactions on chemokine oligomerization. Glycobiology: cwv100. doi:10.1093/glycob/cwv100

11. Eberhardt J, Santos-Martins D, Tillack AF, Forli S. 2021. AutoDock Vina 1.2.0: New docking methods, expanded force field, and python bindings. J Chem Inf Model 61:3891-3898. doi: 10.1021/acs.jcim.1c00203

12. Emsley P, Cowtan K. 2004. Coot: model-building tools for molecular graphics. Acta Crystallographica Section D Biological Crystallography 60:2126–2132. doi:10.1107/S0907444904019158

13. Evans LR, Linker A. 1973. Production and characterization of the slime polysaccharide of *Pseudomonas aeruginosa*. J Bacteriol 116:915–924. doi:10.1128/JB.116.2.915-924.1973

14. Fethiere J, Eggimann B, Cygler M. 1999. Crystal structure of chondroitin AC lyase, a representative of a family of glycosaminoglycan degrading enzymes. J Mol Biol 288:635–647. doi:10.1006/jmbi.1999.2698

15. Garron ML, Cygler M. 2010. Structural and mechanistic classification of uronic acid-containing polysaccharide lyases. Glycobiology 20:1547–1573. doi:10.1093/glycob/cwq122

16. Garron ML, Cygler M. 2014. Uronic polysaccharide degrading enzymes. Curr Opin Struc Biol 28:87–95. doi: 10.1016/j.sbi.2014.07.012

17. Han Y, Garron M, Kim H, Kim W, Zhang Z, Ryu K, Shaya D, Xiao Z, Cheong C, Kim YS, Linhardt RJ, Jeon YH, Cygler M. 2009. Structural snapshots of heparin depolymerization by heparin lyase I. J Biol Chem 284:34019–34027. doi:10.1074/jbc.M109.025338

18. Hashimoto W, Maruyama Y, Nakamichi Y, Mikami B, Murata K. 2014. Crystal structure of *Pedobacter heparinus* heparin lyase Hep III with the active site in a deep cleft. Biochemistry-Us 53:777–786. doi:10.1021/bi4012463

19. Holm L, Rosenstrom P. 2010. Dali server: conservation mapping in 3D. Nucleic Acids Res 38: W545–W549. doi:10.1093/nar/gkq366

20. Huang W, Lunin V, Li Y, Suzuki S, Sugiura N, Miyazono H, Cygler M. 2003. Crystal structure of *Proteus vulgaris* chondroitin sulfate ABC lyase I at 1.9 Å resolution. J Mol Biol 328:623–634. doi:10.1016/S0022-2836(03)00345-0

21. Huang W, Matte A, Li Y, Kim YS, Linhardt RJ, Su H, Cygler M. 1999. Crystal structure of chondroitinase B from *Flavobacterium heparinum* and its complex with a disaccharide product at 1.7 A resolution. J Mol Biol 294:1257- 1269. doi:10.1006/jmbi.1999.3292

22. Irie F, Okuno M, Matsumoto K, Pasquale EB, Yamaguchi Y. 2008. Heparan sulfate regulates ephrin-A3/EphA receptor signaling. P Natl Acad Sci Usa 105:12307–12312. doi:10.1073/pnas.0801302105

23. Itoh T, Nakagawa E, Yoda M, Nakaichi A, Hibi T, Kimoto H. 2019. Structural and biochemical characterisation of a novel alginate lyase from *Paenibacillus* sp. str. FPU-7. Sci Rep-Uk 9. doi:10.1038/s41598-019-51006-1

24. Jagtap SS, Hehemann J, Polz MF, Lee J, Zhao H. 2014. Comparative biochemical characterization of three exolytic oligoalginate lyases from *Vibrio splendidus* reveals complementary substrate scope, temperature, and pH adaptations. Appl Environ Microb 80:4207–4214. doi:10.1128/AEM.01285-14

25. Ji S, Dix SR, Aziz AA, Sedelnikova SE, Baker PJ, Rafferty JB, Bullough PA, Tzokov SB, Agirre J, Li F, Rice DW. 2019. The molecular basis of endolytic activity of a multidomain alginate lyase from *Defluviitalea phaphyphila*, a representative of a new lyase family, PL39. J Biol Chem 294:18077-18091. doi:10.1074/jbc.RA119.010716

26. Kabsch W. 2010. XDS. Acta Crystallogr D Biol Crystallogr 66:125–132. doi:10.1107/S0907444909047337

27. Kusche-Gullberg M, Kjellen L. 2003. Sulfotransferases in glycosaminoglycan biosynthesis. Curr Opin Struc Biol 13:605–611. doi:10.1016/j.sbi.2003.08.002

28. Li S, Jedrzejas MJ. 2001. Hyaluronan binding and degradation by *Streptococcus agalactiae* hyaluronate lyase. J Biol Chem 276:41407–41416. doi:10.1074/jbc.M106634200

29. Lombard V, Bernard T, Rancurel C, Brumer H, Coutinho PM, Henrissat B. 2010. A hierarchical classification of polysaccharide lyases for glycogenomics. Biochem J 432:437–444. doi:10.1042/BJ20101185

30. Lunin VV, Li Y, Linhardt RJ, Miyazono H, Kyogashima M, Kaneko T, Bell AW, Cygler M. 2004. High-resolution crystal structure of *Arthrobacter aurescens* chondroitin AC lyase: an enzyme–substrate complex defines the catalytic mechanism. J Mol Biol 337:367–386. doi:10.1016/j.jmb.2003.12.071

31. Martinez-Fleites C, Smith NL, Turkenburg JP, Black GW, Taylor EJ. 2009. Structures of two truncated phage-tail hyaluronate lyases from *Streptococcus pyogenes serotype* M1. Acta Crystallogr Sect F Struct Biol Cryst Commun 65:963–966. doi:10.1107/S1744309109032813

32. Matsubara Y, Iwasaki K, Muramatsu T. 1998. Action of poly (alpha-L-guluronate) lyase from *Corynebacterium* sp. ALY-1 strain on saturated oligoguluronates. Biosci Biotech Bioch 62:1055-1060. doi:10.1271/bbb.62.1055

33. McCoy AJ, Grosse-Kunstleve RW, Adams PD, Winn MD, Storoni LC, Read RJ. 2007. Phaser crystallographic software. J Appl Crystallogr 40:658–674. doi:10.1107/S0021889807021206

34. Ndeh D, Baslé A, Strahl H, Yates EA, McClurgg UL, Henrissat B, Terrapon N, Cartmell A. 2020. Metabolism of multiple glycosaminoglycans by *Bacteroides thetaiotaomicron* is orchestrated by a versatile core genetic locus. Nat Commun 11 doi:10.1038/s41467-020-14509-4

35. Ochiai A, Yamasaki M, Mikami B, Hashimoto W, Murata K. 2010. Crystal structure of exotype alginate lyase Atu3025 from *Agrobacterium tumefaciens*. J Biol Chem 285:24519–24528. doi:10.1074/jbc.M110.125450

36. Pandey S, Mahanta P, Berger BW, Acharya R. 2021. Structural insights into the mechanism of pH-selective substrate specificity of the polysaccharide lyase Smlt1473. J Biol Chem 297:101014. doi:10.1016/j.jbc.2021.101014

37. Park D, Jagtap S, Nair SK. 2014. Structure of a PL17 family alginate lyase demonstrates functional similarities among exotype depolymerases. J Biol Chem 289:8645-8655. doi:10.1074/jbc.M113.531111

38. Pawar SN, Edgar KJ. 2012. Alginate derivatization: A review of chemistry, properties and applications. Biomaterials 33:3279–3305. doi:10.1016/j.biomaterials.2012.01.007

39. Pettersen EF, Goddard TD, Huang CC, Meng EC, Couch GS, Croll TI, Morris JH, Ferrin TE. 2021. UCSF ChimeraX: Structure visualization for researchers, educators, and developers. Protein Sci 30:70–82. doi:10.1002/pro.3943

40. Popper ZA, Michel G, Hervé C, Domozych DS, Willats WGT, Tuohy MG, Kloareg B, Stengel DB. 2011. Evolution and diversity of plant cell walls: from algae to flowering plants. Annu Rev Plant Biol 62:567–590. doi:10.1146/annurev-arplant-042110-103809

41. Ronne ME, Tandrup T, Madsen M, Hunt CJ, Myers PN, Moll JM, Holck J, Brix S, Strube ML, Aachmann FL, Wilkens C, Svensson B. 2023. Three alginate lyases provide a new gut *Bacteroides ovatus* isolate with the ability to grow on alginate. Appl Environ Microb 89:e0118523. doi:10.1128/aem.01185-23

42. Sattelle BM, Shakeri J, Roberts IS, Almond A. 2010. A 3D-structural model of unsulfated chondroitin from high-field NMR: 4-sulfation has little effect on backbone conformation. Carbohyd Res 345:291–302. doi:10.1016/j.carres.2009.11.013

43. Shaya D, Hahn B, Bjerkan TM, Kim WS, Park NY, Sim J, Kim Y, Cygler M. 2008. Composite active site of chondroitin lyase ABC accepting both epimers of uronic acid. Glycobiology 18:270–277. doi:10.1093/glycob/cwn002

44. Shaya D, Hahn BS, Park NY, Sim JS, Kim YS, Cygler M. 2008. Characterization of chondroitin sulfate lyase ABC from *Bacteroides thetaiotaomicron* WAL2926. Biochemistry-Us 47:6650–6661. doi:10.1021/bi800353g

45. Shaya D, Tocilj A, Li Y, Myette J, Venkataraman G, Sasisekharan R, Cygler M. 2006. Crystal structure of heparinase II from *Pedobacter heparinus* and its complex with a disaccharide product. J Biol Chem 281:15525–15535. doi:10.1074/jbc.M512055200

46. Sirko S, von Holst A, Weber A, Wizenmann A, Theocharidis U, Gotz M, Faissner A. 2010. Chondroitin sulfates are required for fibroblast growth factor-2-dependent proliferation and maintenance in neural stem cells and for epidermal growth factor-dependent migration of their progeny. Stem Cells 28:775–787. doi:10.1002/stem.309

47. Stourac J, Vavra O, Kokkonen P, Filipovic J, Pinto G, Brezovsky J, Damborsky J, Bednar D. 2019. Caver Web 1.0: Identification of tunnels and channels in proteins and analysis of ligand transport. Nucleic Acids Research 47: W414-W422. doi: 10.1093/nar/gkz378.

48. Sugiura N, Setoyama Y, Chiba M, Kimata K, Watanabe H. 2011. *Baculovirus envelope* protein ODV-E66 is a novel chondroitinase with distinct substrate specificity. J Biol Chem 286:29026–29034. doi:10.1074/jbc.M111.251157

49. Trott O, Olson AJ. 2010. AutoDock Vina: improving the speed and accuracy of docking with a new scoring function, efficient optimization, and multithreading. J Comput Chem 31:455-461. doi:10.1002/jcc.21334

50. Ulaganathan T, Shi R, Yao D, Gu R, Garron M, Cherney M, Tieleman DP, Sterner E, Li G, Li L, Linhardt RJ, Cygler M. 2017. Conformational flexibility of PL12 family heparinases: structure and substrate specificity of heparinase III from Bacteroides thetaiotaomicron (BT4657). Glycobiology (Oxford) 27:176-187. doi:10.1093/glycob/cww096

51. Wei L, Zou R, Du M, Zhang Q, Lu D, Xu Y, Xu X, Wang W, Zhang YZ, Li F. 2024. Discovery of a class of glycosaminoglycan lyases with ultrabroad substrate spectrum and their substrate structure preferences. J Biol Chem 300:107466. doi:10.1016/j.jbc.2024.107466

52. Winter WT, Smith PJ, Arnott S. 1975. Hyaluronic acid: structure of a fully extended 3-fold helical sodium salt and comparison with the less extended 4-fold helical forms. J Mol Biol 99:219–235. doi:10.1016/s0022-2836(75)80142-2

53. Yamasaki M, Moriwaki S, Miyake O, Hashimoto W, Murata K, Mikami B. 2004. Structure and function of a hypothetical *Pseudomonas aeruginosa* protein PA1167 classified into family PL-7. J Biol Chem 279:31863–31872. doi:10.1074/jbc.M402466200

54. Yamasaki M, Ogura K, Moriwaki S, Hashimoto W, Murata K, Mikami B. 2005. Crystallization and preliminary X-ray analysis of alginate lyases A1-II and A1-II′ from *Sphingomonas* sp. A1. Acta Crystallogr F 61:288-290. doi:10.1107/S174430910500299X

55. Zhang F, Zheng L, Cheng S, Peng Y, Fu L, Zhang X, Linhardt R. 2019. Comparison of the interactions of different growth factors and glycosaminoglycans. Molecules 24:3360. doi:10.3390/molecules24183360

56. Zhang Q, Cao H, Wei L, Lu D, Du M, Yuan M, Shi D, Chen X, Wang P, Chen X, Chi L, Zhang Y, Li F. 2021. Discovery of exolytic heparinases and their catalytic mechanism and potential application. Nat Commun 12 doi:10.1038/s41467-021-21441-8

57. Zhu B, Huang L, Tan H, Qin Y, Du Y, Yin H. 2015. Characterization of a new endo-type polyM-specific alginate lyase from Pseudomonas sp. Biotechnol Lett 37:409–415. doi:10.1007/s10529-014-1685-0

