## Supplementary data for "Crystal structure and catalytic mechanism of PL35 family glycosaminoglycan lyases with an ultrabroad substrate spectrum"

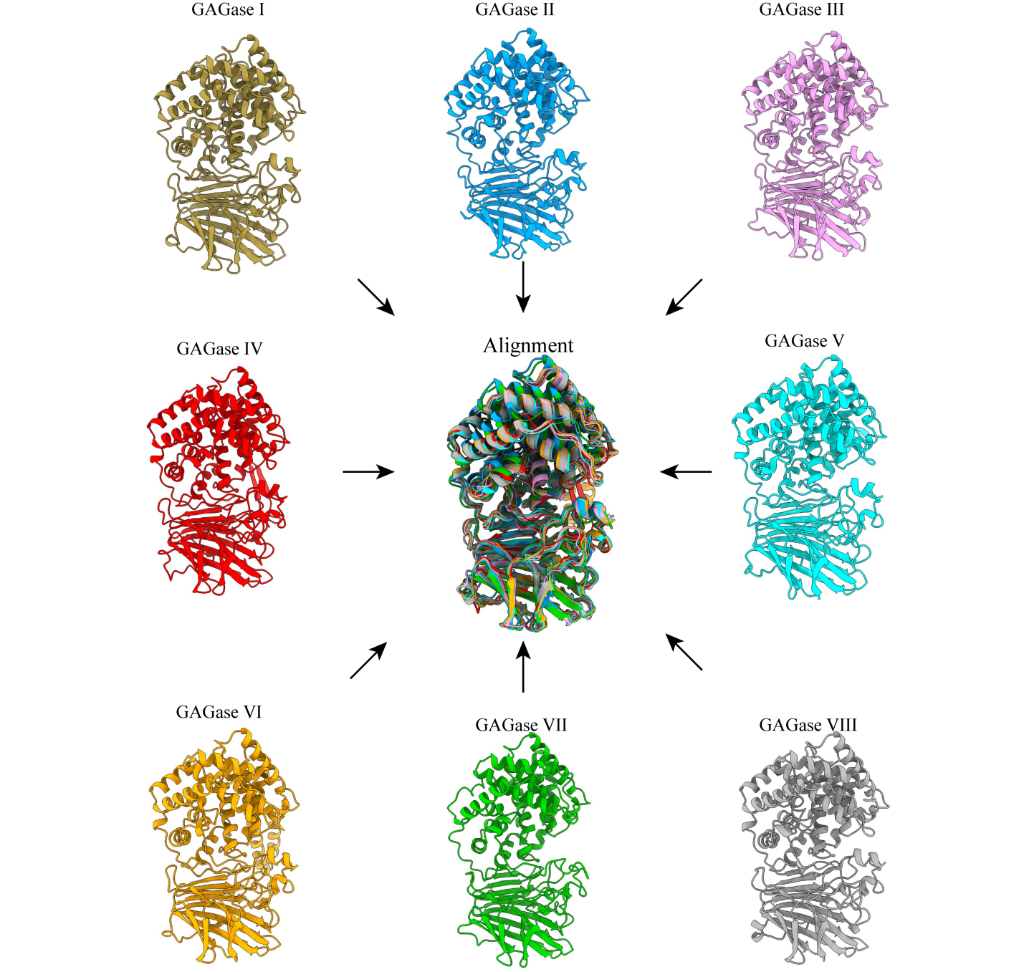

**Figure S1. Structural modeling and alignment of other GAGases.** The structures of the other GAGases were predicted by RoseTTAfold. All of them show good superposition in their predicted structures.

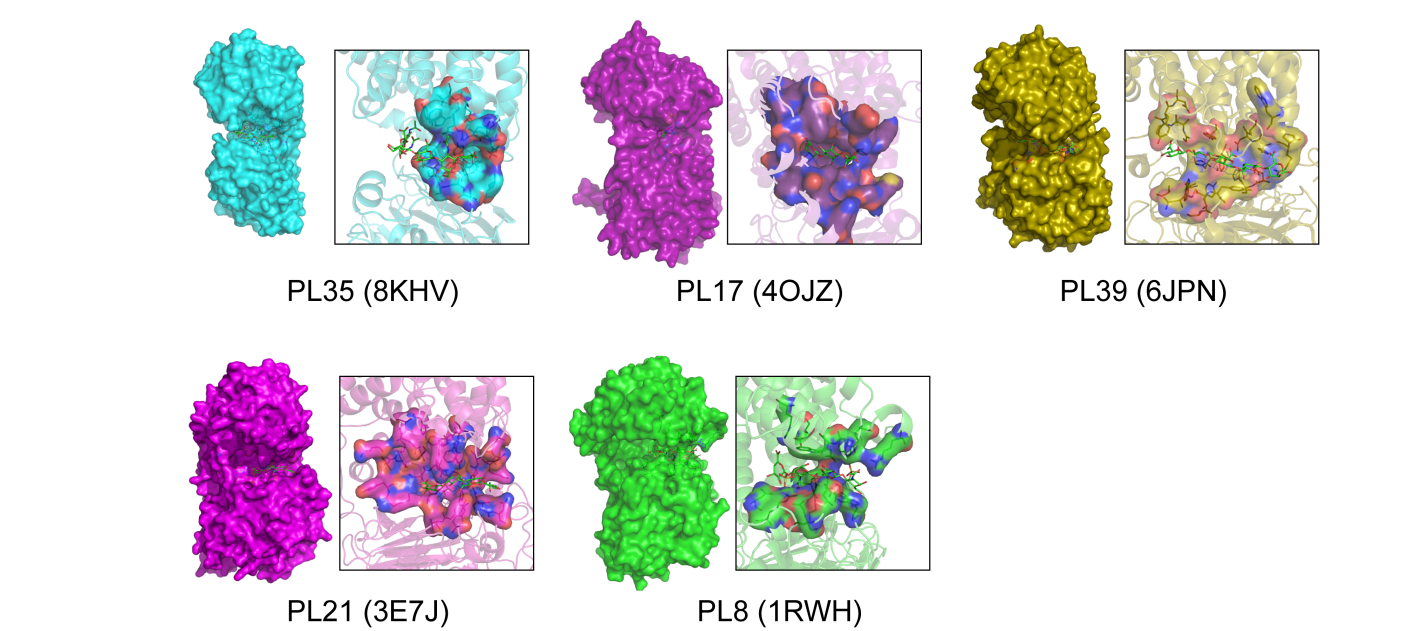

**Figure S2. Surface representations and substrate binding tunnels of GAGase II and its structurally similar GAG/alginate lyase.** The surface representation of GAGase II (8KHV, *blue*) from PL35 family, alginate lyase from PL17 family (4OJZ, *purple*, ligand: β-D-mannopyranuronic acid-(1-4)-α-D-mannopyranuronic acid-(1-4)-α-L-gulopyranuronic acid), alginate lyase from PL39 family (6JPN, *olive*, ligand: β-D-mannopyranuronic acid-(1-4)-β-D-mannopyranuronic acid-(1-4)-β-D-mannopyranuronic acid-(1-4)- β-D-mannopyranuronic acid-(1-4)-β-D-mannopyranuronic acid), heparinase II from PL21 family (3E7J, *magenta*, ligand: 4-deoxy-α-L-threo-hex-4-enopyranuronic acid-(1-4)-2-acetamido-2-deoxy-β-D-glucopyranose-(1-4)-α-D-glucopyranuronic acid-(1-4)-2-acetamido-2-deoxy-β-D-glucopyranose) and chondroitinase AC II from PL8 family (1RWH, *green*, ligand: 6-anhydro-3-deoxy-L-threo-hex-2-enonic acid-(1-3)-2-acetamido-2-deoxy-4-*O*-sulfo-β-D-galactopyranose-(1-4)-2,6-anhydro-3-deoxy-L-xylo-hexonic acid-(1-3)-2-acetamido-2-deoxy-4-*O*-sulfo-β-D-galactopyranose). The dimensions of their **s**ubstrate-binding cavities are measured as follows:14.24 Å (PL35, 8KVI, Asp233-OD2 to Tyr246-OH), 24.25 Å (PL17, 4OJZ, Lys94-NZ to Gln138-NE2), 22.97 Å (PL39, 6JPN, Arg183-NH2 to Tyr343-OH), 25.55 Å (PL21, 3E7J, Glu136-ZN to Lys197-ZN) and 16.97 Å (PL8, 1RWH, Arg134-NH2 to Arg174-NH1).

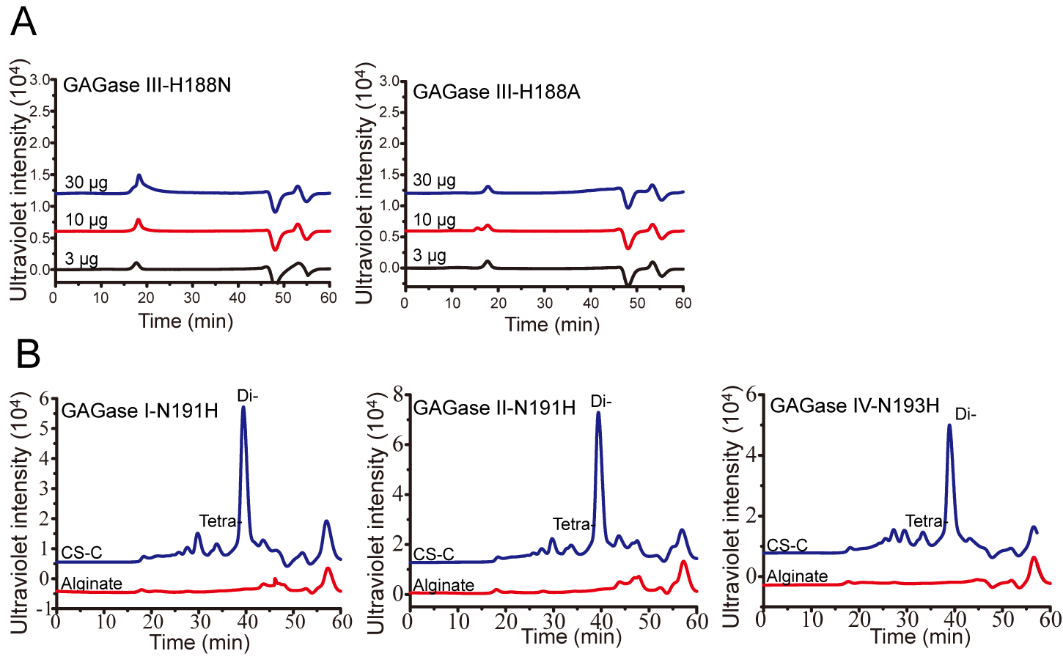

**Figure S3. Activity assay of GAGases variants against CS-C and alginate.** (A), Activity assay of different concentrations of GAGase III-H188N and GAGase III-H188A against alginate; (B), Activity assay of GAGase I-N191H, GAGase II-N191H and GAGase IV-N193H, where the crucial asparagine residues in GAGase I, II and IV were mutated to histidine, respectively. The activities of the corresponding mutants against CS-C or alginate were evaluated using gel filtration HPLC on a Superdex Peptide column as described under “*Materials and methods*”; Di- is indicated as CS disaccharide; Tetra- is indicated as CS tetrasaccharide.

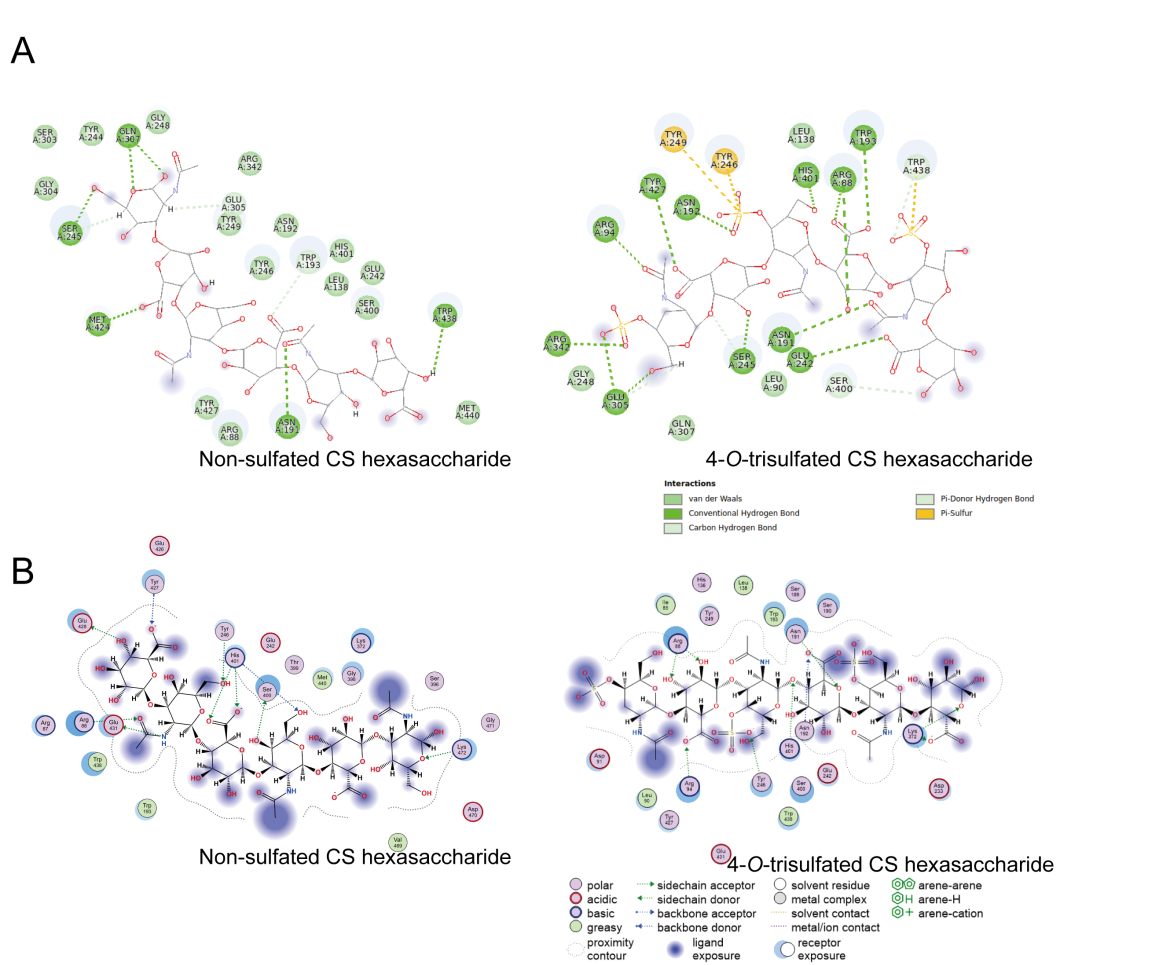

**Figure S4. Two-dimensional interaction plot of GAGase II with molecular docking substrates.** (A), GAGase II form hydrogen bonds, carbon-hydrogen bonds, Pi hydrogen bonds, Pi-Sulfur and van der Waals interactions with nonsulfated (PDB code: 2KQO) (*left*) and 4-*O*-trisulfated (PDB code: 1C4S) (*right*) CS hexasaccharides. Interactions were analysed by discovery studio software. (B), Interaction between GAGase II and nonsulfated CS hexasaccharide (PDB code: 2KQO) (*left*) or 4-*O*-trisulfated CS hexasaccharide (PDB code: 1C4S) (*right*) was analysed using Molecular Operating Environment (MOE).

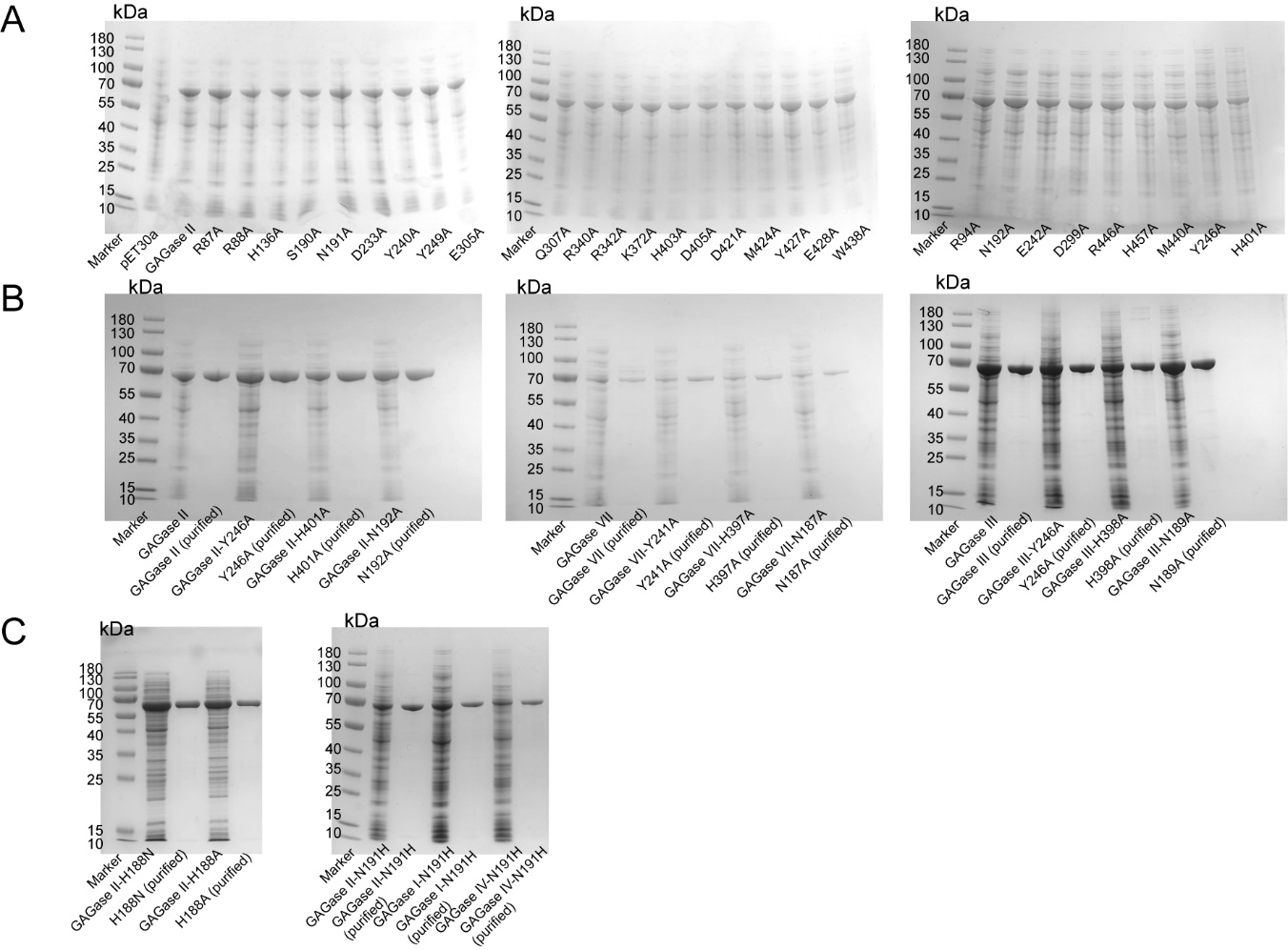

**Figure S5. SDS-PAGE of GAGases and their variants.** The expression and purification of the indicated GAGases and their various mutants, including the putative substrate binding site variants of GAGase II (A), the key triplet residues variants of GAGase II/III/VII (B), and other related variants (C), were assessed by SDS-polyacrylamide gel electrophoresis (SDS-PAGE) followed by staining with Coomassie Brilliant Blue R-250.

**Table S1. Inductively coupled plasma-mass spectrometry (ICP-MS) analysis of GAGase II and GAGase VII.**

|  | Enzyme concentration | Ions  concentration | | | |
| --- | --- | --- | --- | --- | --- |
|  |  | Mn | Ca | Zn | Cu |
|  | (mg/ml) | (μg/L) | | | |
| Negative | - | 0.24 | 24.37 | 6.36 | 0.68 |
| GAGase II | 4.75 | 15.62 | 20.59 | 6.71 | 1.27 |
| GAGase VII | 2.70 | 21.58 | 24.345 | 9.84 | 1.25 |

**Table S2. Sequence information of the identified GAGases.**

|  | GenBank accession number | Originate | Sequence identity* (Query cover/Per. Ident) |
| --- | --- | --- | --- |
| GAGase II | SOD82962.1 | *Spirosoma fluviale* | 100%/100% |
| GAGase VII | EDV05210.1 | *Bacteroides intestinalis* DSM 17393 | 93%/44.7% |
| GAGase I | ADB38475.1 | *Spirosoma linguale* DSM 74 | 100%/81.21 |
| GAGase III | MBC7922758.1 | *Ferruginibacter* sp. | 98%/64.1% |
| GAGase IV | MPR36080.1 | *Prolixibacteraceae bacterium* | 99%/54.6% |
| GAGase V | HCY41582.1 | *Cytophagaceae bacterium* SJW1-29 | 99%/54.0% |
| GAGase VI | EKX96148.1 | *Prevotella saccharolytica* F0055 | 98%/49.1% |
| GAGase VIII | AVM53611.1 | *Bacteroides zoogleoformans* | 98%/42.7% |

*Sequence identity means the sequence similarity compared with GAGase II.

**Table S3. Strains and primers used in this study.**

|  | Description | | Source |
| --- | --- | --- | --- |
| Strain | |  |  |
| *Bacteroides intestinalis* DSM 17393  *E. coli* BL21(DE3) | | Intestinal microorganisms isolated from human faces  F^-^ *omp*T *hsd*S (rB^-^, mB^-^) *gal dcm* (DE3) | DSMZ |
|  |  |  | Vazyme Biotech. |
| Mutant primers | |  |  |
| GAGase II-Y246A-F | | 5’-TATAGCGCGTGGGGCTATGGCACGAGCTTTAA-3’ | Sangon Biotech |
| GAGase II-Y246A-R | | 5’-TAGCCCCACGCGCTATAGCCTTCCGGATACGC-3’ | . |
| GAGase II-H401A-F | | 5’-GCGGCGCACATGGATGTGGGCAGCTTTGTGAT-3’ |  |
| GAGase II-H401A-R | | 5’-ACATCCATGTGCGCCGCGCTCGTGCCCGGGCTGCC-3’ |  |
| GAGase II-N192A-F | | 5’-AACGCGTGGAACCAAGTGTGCAACGCGGGCAT-3’ |  |
| GAGase II-N192A-R | | 5’-ACTTGGTTCCACGCGTTGCTGCTGCGCAGCCA-3’ |  |
| GAGase III-Y243A-F | | 5’-TATGGCGCGTGGGGCTATGGCACGAGCTTTAA-3’ |  |
| GAGase III-Y243A-R | | 5’- TAGCCCCACGCGCCATAGCCTTCCGGATACGC-3’ |  |
| GAGase III-H398A-F | | 5’- GTGAACGCGGCGCACATGGATGTGGGCAGCTT |  |
| GAGase III-H398A-R | | 5’- ATGTGCGCCGCGTTCACGCTCGGGCT-3’ |  |
| GAGase III-N189A-F | | 5’-CATGCGTGGAACCAAGTGTGCAACG-3’ |  |
| GAGase III-N189A-R | | 5’-ACTTGGTTCCACGCATGGCTCGCTTTCAGCC-3’ |  |
| GAGase VII-Y241A-F | | 5’-TATTCCGCGTGGGGATATGGGACGACTTACAA-3’ |  |
| GAGase VII-Y241A-R | | 5’-TATCCCCACGCGGAATAGCCTTCGGCATACGC-3’ |  |
| GAGase VII-H397A-F | | 5’-ATCAGGAGCGACACATTTGGATGCCGGTTCTT-3’ |  |
| GAGase VII-H397A-R | | 5’-AATGTGTCGCTCCTGATTGTGCTGTCCCTCCT-3’ |  |
| GAGase VII-N187A-F | | 5’-TAATGCGTGGAATCAGGTATGTAATGGGGGAA-3’ |  |
| GAGase VII-N187A-R | | 5’-CCTGATTCCACGCATTATTTCGATAGAGCCAGCTATTATATT-3’ |  |
| GAGase II-R87A-F | | 5’-TATTCAGATTGGCGCGCGCCTGCTGGATAAAAGCC-3’ |  |
| GAGase II-R87A-R | | 5’-GCGCGCCAATCTGAATACGTTTCAGCGGTTCA-3’ |  |
| GAGase II-R88A-F | | 5’-CGCGCTGCTGGATAAAAGCCGCGAAGCGCTGC-3’ |  |
| GAGase II-R88A-R | | 5’-TTTATCCAGCAGCGCGCGGCCAATCTGAATACGTT-3’ |  |
| GAGase II-R94A-F | | 5’-ATAAAAGCGCGGAAGCGCTGCGCCGCATTTTT-3’ |  |
| GAGase II-R94A-R | | 5’-CGCTTCCGCGCTTTTATCCAGCAGGCGGCGGC-3’ |  |
| GAGase II-H136A-F | | 5’-ACCGCGTTTCTGGATGTGGCGGAAATGACCAT-3’ |  |
| GAGase II-H136A-R | | 5’-ACATCCAGAAACGCGGTCGGGTTCCAATCGCT-3’ |  |
| GAGase II-N191A-F | | 5’-GCGAACTGGAACCAAGTGTGCAACGCGGGCAT-3’ |  |
| GAGase II-N191A-R | | 5’-ACTTGGTTCCAGTTCGCGCTGCTGCGCAGCCAGCT-3’ |  |
| GAGase II-D233A-F | | 5’-GATGGGCGCGTATAAACCGGATGGCGCGTATC-3’ |  |
| GAGase II-D233A-R | | 5’-GTTTATACGCGCCCATCGGCAGCACCACCGCG-3’ |  |
| GAGase II-Y240A-F | | 5’-GGCGCCGGAAGGCTATAGCTATTGGGGCTATG-3’ |  |
| GAGase II-Y240A-R | | 5’-TATAGCCTTCCGGCGCCGCGCCATCCGGTTTATA-3’ |  |
| GAGase II-E242A-F | | 5’-GGCGGGCTATAGCTATTGGGGCTATGGCACGA-3’ |  |
| GAGase II-E242A-R | | 5’-AATAGCTATAGCCCGCCGGATACGCGCCATCCGG-3’ |  |
| GAGase II-Y249A-F | | 5’-ATAGCTATTGGGGCGCGGGCACGAGCTTTAACGTGATG-3’ |  |
| GAGase II-Y249A-R | | 5’-CGCGCCCCAATAGCTATAGCCTTCCGGATACG-3’ |  |
| GAGase II-D299A-F | | 5’-TAACTATAGCGCGAGCGGCCTGAGCGGCGAAC-3’ |  |
| GAGase II-D299A-R | | 5’-CGCTCGCGCTATAGTTATACGCGTTGCCGCTC-3’ |  |
| GAGase II-Q307A-F | | 5’-TGGCGCCGGCGATGTTTTGGTTTGCGAAAAAA-3’ |  |
| GAGase II-Q307A-R | | 5’-AAACATCGCCGGCGCCAGTTCGCCGCTCAGGCC-3’ |  |
| GAGase II-R340A-F | | 5’-AGAACCATCTGGCGAACCGCCTGCTGCCGGCG-3’ |  |
| GAGase II-R340A-R | | 5’-GTTCGCCAGATGGTTCTGCGGGTTGCTGTTCA-3’ |  |
| GAGase II-R342A-F | | 5’-ATCTGCGCAACGCCCTGCTGCCGGCGGCGCTG-3’ |  |
| GAGase II-R342A-R | | 5’-CAGGGCGTTGCGCAGATGGTTCTGCGGGTTGC-3’ |  |
| GAGase II-H403A-F | | 5’-CATGCGGCGATGGATGTGGGCAGCTTTGTGAT-3’ |  |
| GAGase II-H403A-R | | 5’-ACATCCATCGCCGCATGGCTCGTGCCCGGGCT-3’ |  |
| GAGase II-D405A-F | | 5’-ATGCGCACATGGCGGTGGGCAGCTTTGTGATGGA-3’ |  |
| GAGase II-D405A-R | | 5’-CACCGCCATGTGCGCATGGCTCGTGCCCGGGC-3’ |  |
| GAGase II-D421A-F | | 5’-ATGGCGTTTGGCATGCAAGAATATGAAAGCCT-3’ |  |
| GAGase II-D421A-R | | 5’-TGCATGCCAAACGCCATCGCCCAGCGCACGCC-3’ |  |
| GAGase II-M424A-F | | 5’-TTTTGGCGCGCAAGAATATGAAAGCCTGGAAAGC-3’ |  |
| GAGase II-M424A-R | | 5’-ATTCTTGCGCGCCAAAATCCATCGCCCAGCGC-3’ |  |
| GAGase II-Y427A-F | | 5’-TGCAAGAAGCGGAAAGCCTGGAAAGCAAAGGC-3’ |  |
| GAGase II-Y427A-R | | 5’-GCTTTCCGCTTCTTGCATGCCAAAATCCATCG-3’ |  |
| GAGase II-E428A-F | | 5’-AGAATATGCGAGCCTGGAAAGCAAAGGCGTGG-3’ |  |
| GAGase II-E428A-R | | 5’-CCAGGCTCGCATATTCTTGCATGCCAAAATCCA-3’ |  |
| GAGase II-E431A-F | | 5’-AAAGCCTGGCGAGCAAAGGCGTGGATCTGTGG-3’ |  |
| GAGase II-E431A-R | | 5’-TTTGCTCGCCAGGCTTTCATATTCTTGCATGC-3’ |  |
| GAGase II-W438A-F | | 5’-GGATCTGGCGAACATGAAACAGAACAGTCAGCGC-3’ |  |
| GAGase II-W438A-R | | 5’-TCATGTTCGCCAGATCCACGCCTTTGCTTTCC-3’ |  |
| GAGase II-M440A-F | | 5’-GTGGAACGCGAAACAGAACAGTCAGCGCTGGC-3’ |  |
| GAGase II-M440A-R | | 5’-TCTGTTTCGCGTTCCACAGATCCACGCCTTTG-3’ |  |
| GAGase II-R446A-F | | 5’-ACAGAACAGTCAGGCGTGGCAGATTCTGCGCTATAACA-3’ |  |
| GAGase II-R446A-R | | 5’-ACGCCTGACTGTTCTGTTTCATGTTCCACAGA-3’ |  |
| GAGase II-H457A-F | | 5’-ACTTTGCGGCGAACACCCTGAGCATTAACGATGA-3’ |  |
| GAGase II-H457A-R | | 5’-GTGTTCGCCGCAAAGTTGTTATAGCGCAGAAT-3’ |  |
| GAGase III-H188A-F | | GCGAACTGGAACCAAGTGTGCAACGCGGGCAT |  |
| GAGase III-H188A-R | | ACTTGGTTCCAGTTCGCGCTCGCTTTCAGCCAGTTGT |  |
| GAGase III-H188N-F | | AGCGAGCAATAACTGGAACCAAGTGTGCAACG |  |
| GAGase III-H188N-R | | TCCAGTTATTGCTCGCTTTCAGCCAGTTGTTA |  |
| GAGase II N191H-F | | AGCAGCCACAACTGGAACCAAGTGTGCAACGC |  |
| GAGase II N191H-R | | TTCCAGTTGTGGCTGCTGCGCAGCCAGCTGTT |  |
